# The evolutionary conserved complex CEP90, FOPNL and OFD1 specifies the future location of centriolar distal appendages, and promotes their assembly

**DOI:** 10.1101/2021.07.13.452210

**Authors:** Pierrick Le Borgne, Logan Greibill, Marine Hélène Laporte, Michel Lemullois, Khaled Bouhouche, Mebarek Temagoult, Olivier Rosnet, Maeva Le Guennec, Laurent Lignières, Guillaume Chevreux, France Koll, Virginie Hamel, Paul Guichard, Anne-Marie Tassin

**Affiliations:** Université Paris-Saclay, CEA, CNRS, Institute for Integrative Biology of the Cell (I2BC), 91198, Gif-sur-Yvette, France; University of Geneva, Department of Cell Biology, Geneva, Switzerland; Imagerie-Gif Light facility, Université Paris-Saclay, CEA, CNRS, Institute for Integrative Biology of the Cell (I2BC), 91198, Gif-sur-Yvette, France; Centre de Recherche en Cancérologie de Marseille (CRCM), Aix Marseille Univ, CNRS, INSERM, Institut Paoli-Calmettes, 13009 Marseille, France; ProteoSeine@IJM, Université de Paris/CNRS, Institut Jacques Monod, 75006 Paris, France

## Abstract

In metazoa, cilia assembly is a cellular process that starts with centriole to basal body maturation, migration to the cell surface and docking to the plasma membrane. Basal body docking involves the interaction of both the distal end of the basal body and the transition fibers / distal appendages, with the plasma membrane. Mutations in numerous genes involved in basal body docking and transition zone assembly are associated with the most severe ciliopathies, highlighting the importance of these events in cilium biogenesis. In this context, the ciliate *Paramecium* has been widely used as a model system to study basal body and cilia assembly. However, despite the apparent evolutionary conservation of cilia assembly events across phyla, whether the same molecular players are functionally conserved, is not fully known. Here, we demonstrated that CEP90, FOPNL and OFD1 form an evolutionary conserved complex that is crucial for ciliogenesis. Using ultrastructure expansion microscopy, we unveiled that these proteins localize at the distal end of both centrioles/basal bodies in *Paramecium* and mammalian cells. Moreover, we found that these proteins are recruited early after centriole duplication on the external surface of the procentriole and define the future location of the distal appendages. Functional analysis performed both in *Paramecium* and mammalian cells demonstrate the requirement of this complex for distal appendage assembly and basal body docking. Finally, we show that mammals require another component, Moonraker (MNR), to recruit OFD1, FOPNL, and CEP90, which will then recruits the distal appendage protein CEP83. Altogether, we propose that this ternary complex is required to determine the future position of distal appendages.

## Introduction

The centrosome is the main microtubule-organizing center in most animal cells. It is composed of two centrioles, microtubule based organelles displaying a nine-fold symmetry, surrounded by layers of pericentriolar material (Paintrand *et al*, 1992; Tassin & Bornens, 1999; Mennella *et al*, 2014). A structural asymmetry between the two centrioles is observed, the mother one bearing distal and subdistal appendages, while the daughter lacks these appendages (Paintrand *et al*, 1992). In resting cells, the mother centriole converts into a basal body (BB), which functions as a platform to template the assembly of a cilium. Cilia can be found at the cell surface either unique or in multiple copies, motile or non-motile (Mitchison & Valente, 2017). Although well conserved throughout the evolution (Vincensini *et al*, 2011), cilia ultrastructure exhibits variations between cell types in a given organism and between organisms (Sorokin, 1962; Garcia-Gonzalo & Reiter, 2017; Wiegering *et al*, 2018; Vieillard *et al*, 2016). In all cases, cilia possess mechano- and chemo-sensitivity properties. In addition, motile cilia beat to generate either a flow such as cerebral fluid (brain ventricle), mucus (respiratory tract) or the forces required to propulse the cells (spermatozoid, ovum, unicellular organisms) (Mitchison & Valente, 2017).

The cilia assembly process, also called ciliogenesis, is a multistep process involving 4 major events: BB duplication, migration to the cell surface, membrane anchoring of the BB via the distal appendages, and ciliary growth. The conservation of this sequence of events in most phyla is paralleled by an important conservation of the proteins involved (Vincensini *et al*, 2011; Jana *et al*, 2014). The BB anchoring step requires the tethering of the distal appendages, which in mammalian cells contain CEP83, CEP89, SCLT1, FBF1 and CEP164 (Tanos *et al*, 2013; Joo *et al*, 2013; Failler *et al*, 2014; Yang *et al*, 2018), to a membrane. This membrane could be either Golgi-apparatus derived-vesicles that fuse together to form the ciliary shaft, as in most metazoan cells, or the plasma membrane directly as in the unicellular organisms *Paramecium* and in some mammalian cell types such as the immunological synapse (Stinchcombe *et al*, 2015). The interaction of the BB with the membrane leads to the formation of the transition zone (TZ), bridging the BB to the axoneme, which has recently been recognized to act as a diffusion barrier between the intracellular space and the cilium, defining the ciliary compartment (Reiter *et al*, 2012; Gonçalves & Pelletier, 2017). Mutations in genes encoding proteins localized at the BB distal end, distal appendages or the TZ lead to various syndromes in humans, called ciliopathies (Oro-facial-digital, Nephronophtysis, Joubert, Jeune), reinforcing the importance to understand how these structures and their associated proteins control ciliogenesis (Singla *et al*, 2010; Lopes *et al*, 2011a; Thauvin-Robinet *et al*, 2014; Failler *et al*, 2014; Chevrier *et al*, 2016).

*Paramecium tetraurelia* is a free-living unicellular organism, easy to cultivate that bear at its surface ca. 4000 cilia. The corresponding BB are arranged in longitudinal rows, and their polarity is marked by the asymmetrical organization of their associated appendages (Iftode *et al*, 1989; Tassin *et al*, 2015). In *Paramecium*, BB organization is highly precise, with regions composed of units with singlet or doublet BB. Units with doublets BB can display either 2 ciliated BB or a ciliated and an unciliated one (Iftode *et al*, 1989). Interestingly, the TZ matures biochemically and structurally between the two states (Gogendeau *et al*, 2020). Unlike metazoa, there is no centriolar stage in *Paramecium:* new BB develop from the docked ones. Once duplicated, they tilt-up and anchor directly at the surface. Due to the high number of BB at the cell surface, a defect in BB anchoring is easily detected in *Paramecium* by immunofluorescence, making of *Paramecium* an outstanding model to characterize new partners involved in BB anchoring.

The centriolar protein FOPNL/FOR20 together with OFD1 belong to the family of FOP-related proteins, displaying a TOF and a LisH domains; the latter one is known to be involved in homodimer formation (Azimzadeh *et al*, 2008). Previous work highlighted a role for FOPNL in ciliogenesis, since its depletion in mammalian cells inhibits cilia formation (Sedjaï *et al*, 2010) and FOPNL KO mice show embryonic lethality at about E11.5 with an important decrease of cilia in the embryonic node. As a consequence, the embryos show left-right symmetry defects (Xu *et al*, 2021). However, the dual localization of FOPNL, at both centrioles and centriolar satellites prevented determining which pool of FOPNL is involved in ciliogenesis, since centriolar satellites are also involved in cilia formation (Sedjaï *et al*, 2010; Stowe *et al*, 2012). Interestingly, the depletion of the FOPNL ortholog in the unicellular organism *Paramecium* led to the inhibition of BB distal end assembly and, as a consequence, blocks the BB docking process (Aubusson-Fleury *et al*, 2012). Since *Paramecium* does not have centriolar satellites as determined by the absence of both PCM1 and granular structures (personal observations) hinting for a direct role of FOPNL at the level of centriole rather than at centriolar satellite. Further studies in human cells strengthened this hypothesis since FOPNL was shown to interact directly with MNR/OFIP/KIAA0753 in a complex comprising the distal centriolar protein OFD1; the presence of FOPNL increasing the interaction between MNR and OFD1 (Chevrier *et al*, 2016). In mammals, OFD1 controls centriole length and is required for distal appendage formation, while in *Paramecium* it is necessary for distal end assembly and BB docking similarly to FOPNL (Singla *et al*, 2010). Recently, Kumar and colleagues shed light on this process by identifying the DISCO complex composed of OFD1, MNR and CEP90 as important for distal appendages formation and proper ciliogenesis (Kumar *et al*, 2021). Despite these convergent information, the role of FOPNL at centrioles and its relationship with other members of the centriolar distal complex such as DISCO remained unknown.

Here, in order to fully characterize the function of the evolutionary conserved protein FOPNL in BB docking and distal appendages assembly, we undertook the identification of proteins in proximity of FOPNL/FOR20 by BioID in human cells. We focused our study on CEP90/PIBF1, since its encoding gene was found mutated in some ciliopathies (Wheway *et al*, 2015; Shen *et al*, 2020). Moreover, CEP90 is evolutionarily conserved from protists to Humans, as OFD1 and FOPNL, despite being absent in some groups as drosophilidae or rabditidae. We demonstrated, through functional analysis performed both in *Paramecium* and mammalian cells using RNAi (RNA interference), the involvement of CEP90 in BB anchoring as well as unveiled the interdependent recruitment of CEP90 with OFD1 and FOPNL/FOR20 to build the distal end of the BB in *Paramecium*. Furthermore, we uncovered, using ultrastructure expansion microscopy (U-ExM), that CEP90, OFD1 and FOPNL localize in *Paramecium* at the distal end of BB in a 9-fold symmetry. Similarly and consistently to the recent work (Kumar *et al*, 2021), we confirmed in mammalian cells that CEP90 and OFD1 localize sub-distally in a 9-fold symmetry as soon as procentriole formation starts and are required, with FOPNL, to recruit distal appendage proteins. Next, we took advantage of a microtubule displacement assay, whereby overexpressed MNR relocates to cytoplasmic microtubules, to analyze protein recruitment and complex formation (Chevrier *et al*, 2016; Kumar *et al*, 2021). Consistently, we found that overexpressed MNR recruits overexpressed OFD1, FOPNL and CEP90 to microtubules, suggesting that they can interact. In addition, MNR and CEP90 co-expression allows the recruitment of endogenous OFD1 as well as the distal appendage protein CEP83 on microtubules. In conclusion, our results demonstrate the evolutionary conserved function of the complex FOPNL, OFD1, CEP90 and MNR in mammalian cells, which predetermines the location of distal appendages and their assembly, allowing BB docking at the cell membrane.

## Results

To discover novel proteins involved in the BB anchoring process, BioID was performed using FOPNL as a bait in human cells. To do so, Flp-In HEK293 cells expressing myc-BirA*-FOPNL under the control of the tetracycline repressor were generated. Centrosomal biotinylated proteins were purified on streptavidin beads and analyzed by mass spectrometry. Candidates were ranked according to their fold change and p-value (Figure S1A). Forty-eight proteins were identified with a fold change >5 and a p-value < 0.05 (Table S1) and two main protein networks were generated using STRING DB to further analyze the data. Consistently, one displayed centriolar satellite proteins as expected from the localization of FOPNL and the other one corresponded to nuclear enriched proteins (Figure S1B), some of them being involved in chromatin organization as previously observed by (Gheiratmand *et al*, 2019). Importantly, several known BB anchoring proteins such as Talpid3 (Kobayashi *et al*, 2014), OFIP/MNR/KIAA0753 (Chevrier *et al*, 2016b), were enriched in our mass spectrometry, thus validating our approach (see Table S1). In addition, some novel potential interactors were found. We focused further our functional analysis on the candidate centriolar protein CEP90/PIBF1 as: i) *CEP90* gene was found mutated in some ciliopathies (Wheway *et al*, 2015); ii) *CEP90* is well conserved from protists to mammals, despite being absent in some phylla.

To validate the putative interaction of FOPNL and CEP90, we first performed co-immunoprecipitation experiments on Hela Kyoto TransgeneOmics cells expressing GFP-tagged mouse FOPNL (Chevrier *et al*, 2016). We found that GFP-FOPNL was able to interact with endogenous OFD1 and CEP90 (Figure S1C).

### Paramecium CEP90-GFP localizes to basal body

To further study the novel interactions found in mammalian cells and to test their evolutionary conservation, we turned to *Paramecium*, a well-established model to decipher the BB anchoring process. We first searched for CEP90 homologs in the *Paramecium tetraurelia* genome and found two CEP90 homologs, *CEP90a* and *CEP90b*, derived from the last whole genome duplication (Aury *et al*, 2006). RNAseq analysis of the *P. tetraurelia* transcriptome during vegetative growth revealed that CEP90a is 24x more expressed than CEP90b (Arnaiz *et al*, 2010, and paramecium.i2bc.paris-saclay.fr), therefore we decided to focus on CEP90a for the rest on this study.

To ascertain the localization of CEP90 in *Paramecium*, we expressed CEP90a-GFP under the control of the constitutive calmodulin gene-regulatory elements. After transformation, rows of GFP dots were observed on transformants displaying a wild type growth rate and phenotype (Figure 1A). To confirm that this localization pattern corresponds to BB, we co-stained with tubulin poly-glutamylation (poly-E), a known BB marker (Gogendeau et al, 2020). Consistently, we found that CEP90a-GFP decorates the BB (Figure 1A) and that this staining is reminiscent to GFP-FOPNL (Figure 1B) and OFD1-GFP staining (Aubusson-Fleury *et al*, 2012; Bengueddach *et al*, 2017), as expected for a proximity partner. A careful observation of the GFP staining pattern relative to the BB, revealed that all BB are decorated, i.e., ciliated and unciliated (Figure 1A, magnification), unlike the GFP-tagged MKS and NPHP protein complex, which stained only ciliated BB (Gogendeau *et al*, 2020). Double labelling of CEP90a-GFP cells using poly-E antibodies and anti-epiplasm (Jeanmaire-Wolf *et al*, 1993), a superficial cytoskeleton layer, revealed that the CEP90a-GFP staining is located at the level of the epiplasmin layer, in which the distal ends of the BB are embedded. This result demonstrates that CEP90 localization is restricted to the distal ends of anchored BB (Figure 1A’, magnification, yellow arrowhead) similarly to GFP-FOPNL (Figure 1B’, magnification, yellow arrowhead).

**Figure 1:**
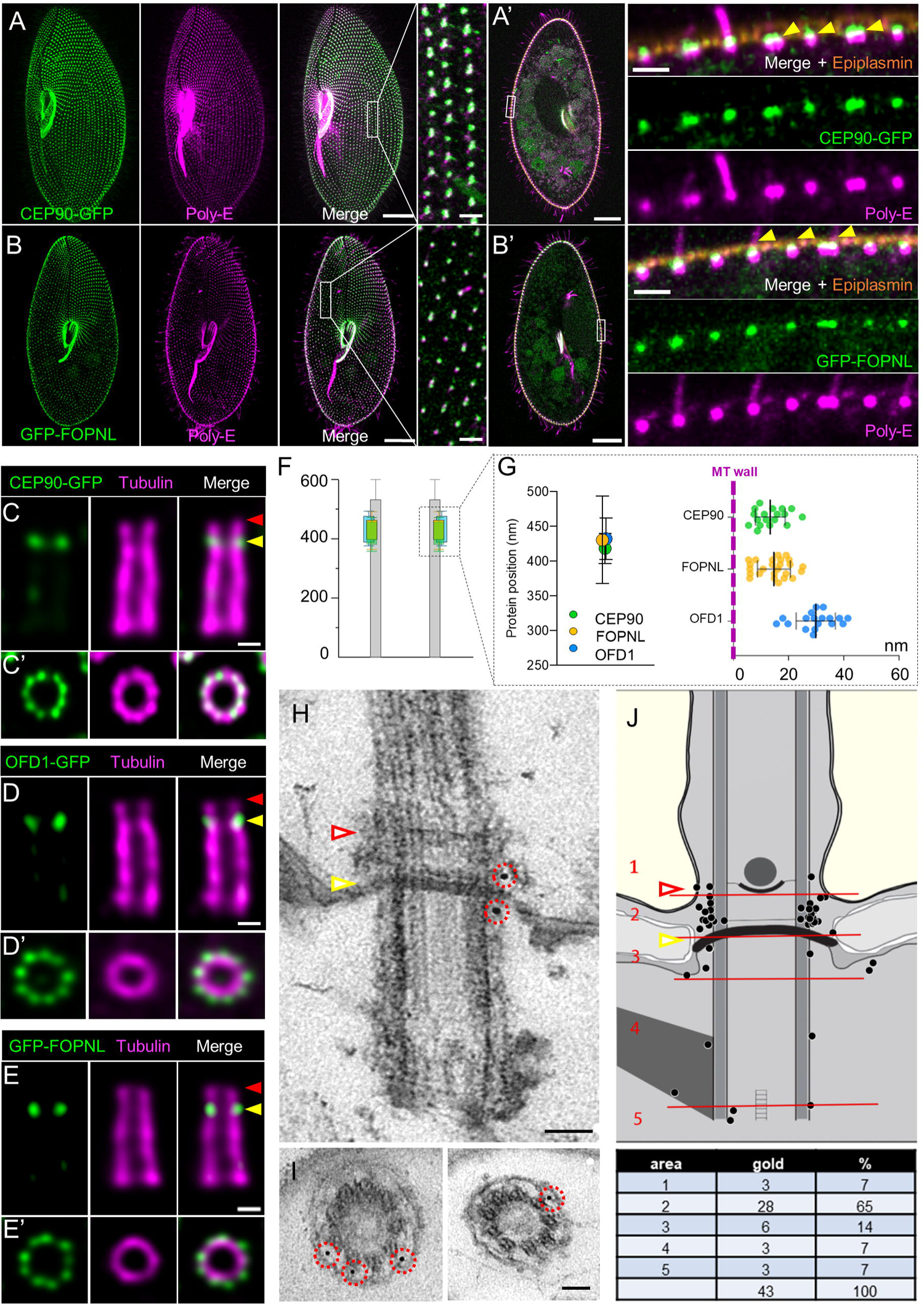
Localization of CEP90 GFP, GFP-FOPNL and OFD1-GFP in *Paramecium*. **A-B’**) Labelling of CEP90-GFP (**A, A’**) and GFP-FOPNL (**B, B’**) paramecia transformants using the polyclonal anti-polyglutamylated tubulin antibodies (Poly-E, magenta). **A, B**) Projection of confocal sections through the ventral surface. On the ventral side (**A, B**), the green emitted fluorescence of GFP overlapped with the anti-poly-E labeling on all BB, as shown on the magnification on the right panel. Scale bars= 20 µm and 2 µm (magnification). **A’, B’**) Projection of confocal sections at the intracytoplasmic level. BB at the cell surface showed that the GFP signal (green) is located at the distal part of all basal bodies (magenta) at the level of the epiplasm layer (orange) for both CEP90-GFP and GFP-FOPNL. The yellow arrowheads indicate the level of the epiplasm layer representing the frontier between the BB and the TZ. Scale bars= 20 µm and 1.7µm (magnification).**C-G**) Cortex from transformants expressing CEP90-GFP, OFD1-GFP and GFP-FOPNL were purified and expanded using the U-ExM technique. Gullet cortical fragments displaying ciliated BB were selected since U-ExM gave better results in this region. Anti-GFP antibodies (green) and anti-tubulin antibodies (magenta) were used to label respectively the fusion protein and the BB microtubule wall. **C-E’**) Labeling of the fusion protein on BB observed longitudinally (upper panel) or top view (lower panel). CEP90-GFP (**C**), OFD1-GFP (**D**) and GFP-FOPNL (**E**) BB display the GFP staining close to the microtubule wall of the BB (magenta) and slightly underneath the distal end staining of the anti-tubulin signal. Yellow and red arrowheads indicate the level of the epiplasm layer and the distal part of the TZ respectively. Top view images demonstrate that the GFP signal is organized in a 9-fold symmetry (**C’, D’, E’**). Scale bar= 100nm. **F, G**) The distance to the proximal end and microtubule wall were quantified and statistically analyzed (***p<0.0001, one-way ANOVA followed by Tukey’s post hoc test). N= 14 (CEP90), 15 (FOPNL), 9 (OFD1) BB from 2 independent experiments. Average +/- SD: CEP90= 418.4 +/- 21.7 nm; FOPNL=430.6 +/- 62.6 nm; OFD1=432.3 +/- 29.9 nm. **H-J**) Representative EM images of the immunolocalization of CEP90-GFP protein on BB revealed by anti-GFP antibodies. **H, I**) longitudinal view (**H**) and transverse sections of BBs at the level of the intermediate plate (**I**). Note that gold particles are located close to the TZ or at the level of the intermediate plate Scale bar= 100nm. **J**) BB diagrams recapitulating the localization of gold beads in the 5 different zones indicated (1-5). Quantification of gold particle number counted on 35 BB in 2 independent experiments. Note that most of the beads were localized at the TZ level between the axosomal plate and the terminal plate, known to be the place where the transition fibers emerge from the microtubule doublet. The yellow and red arrowheads indicate the level of the epiplasm layer and the distal part of the TZ respectively.

### CEP90, FOPNL and OFD1 localizes at the basal body distal ends in a 9-fold symmetry in Paramecium

To characterize more precisely the localization of CEP90 together with FOPNL and OFD1, we turned to U-ExM, a super-resolution method based on the physical expansion of the biological sample in a swellable polymer (Gambarotto *et al*, 2019; Gambarotto *et al*, 2021). Since *Paramecium* is 150 µm long and 50µm thick, U-ExM was performed on isolated cortex sheets (Le Guennec *et al*, 2020) from *Paramecium* expressing GFP-FOPNL, CEP90a-GFP or OFD1-GFP to allow a nanoscale localization of these proteins within the structure (Figure 1C-G). During cortex purification, we observed by electron microscopy (EM) ciliary shedding just above the TZ (Figure S2A, white arrowhead), as it is known to occur in physiological conditions (Adoutte *et al*, 1980; Gogendeau *et al*, 2020). Double stained longitudinally oriented BB show that in CEP90a-GFP paramecia, the GFP staining was located either at the top of the BB assessed by the anti-tubulin staining (Figure S2B, yellow arrowhead) or slightly underneath (Figure S2B, white arrowhead). This observed difference could be associated with the ciliation status, since ciliated BB showed a longer TZ with extended microtubule doublets than unciliated ones (Figure S2A) (Tassin *et al*, 2015b; Gogendeau *et al*, 2020). Therefore, we propose that the staining is found at the proximal part of the TZ (Figure 1C-E, Figure S2 B-E, green arrowhead) on all BB and the observed tubulin staining above the GFP staining might be associated to ciliated BB (Figure 1C-E, Figure S2 B-E, magenta arrowhead) and would correspond to the elongation of the TZ that occurs during ciliogenesis. Longitudinally oriented BB stained for OFD1-GFP (Figure 1D, Figure S2C,E) and GFP-FOPNL (Figure 1E, Figure S2D) BB show a similar pattern than CEP90a-GFP, suggesting that these three proteins localized at the end of the BB or at the proximal part of the TZ (Figure 1C-F). Moreover, analysis of the protein’s positions along the microtubule wall indicated that all three proteins localize at the same average position at around 425 nm from the proximal extremity (Figure 1F, G). Observations on top-viewed BB showed that the three proteins were organized in a 9-fold symmetry, close to the microtubule wall but externally (Figure 1C’, D’, E’). Quantification of the distance between both GFP and tubulin maximal intensity signal of the rings revealed that CEP90a-GFP, OFD1-GFP and GFP-FOPNL localized close to the microtubule wall with CEP90 and FOPNL appearing at around 15 nm away from the microtubule wall, while OFD1 was at around 30 nm, suggesting that OFD1 is more external than the 2 others (Figure 1F, G). This localization was further refined by immunogold staining of CEP90a-GFP on whole permeabilized cells in which the gold particles were found mostly at the level of the terminal/intermediate plates, close to the microtubular wall, where transition fibers are known to emerge in *Paramecium* (Dute & Kung, 1978; Figure 1H-J). Counting of gold particles confirmed that most of the particles are localized at the proximal part of the TZ (65%) (Figure 1J).

### CEP90 depletion prevents basal body docking in Paramecium

Since CEP90 was found at proximity of FOPNL by BioID and localized similarly as OFD1-GFP and GFP-FOPNL, we predicted that it will be involved in BB anchoring through its contribution in building the BB distal end. To test this hypothesis, we first decided to knock-down *CEP90* in *Paramecium* by feeding wild type cells with dsRNA-producing bacteria to induce RNAi silencing (Galvani & Sperling, 2002). As a control, we inactivated the unrelated gene *ND7*, involved in the exocytosis of trichocystes. The high percentage of identity between *CEP90a* and *CEP90b* genes made difficult to design RNAi constructs to inactivate each gene individually, therefore, we silenced them both together. *CEP90* knock-down induced a modification of the cell size and shape from the first division after what they appeared smaller and rounder (Figure 2A, CEP90^RNAi^) until the third/fourth division where they start dying (Figure S3B). The swimming velocity was also reduced, as previously observed for defective BB (Figure S3C, (Gogendeau *et al*, 2020)). The efficiency of the *CEP90* silencing was tested by immunofluorescence by verifying the effective depletion of GFP-CEP90a (Figure 2B, Figure S3A).

**Figure 2:**
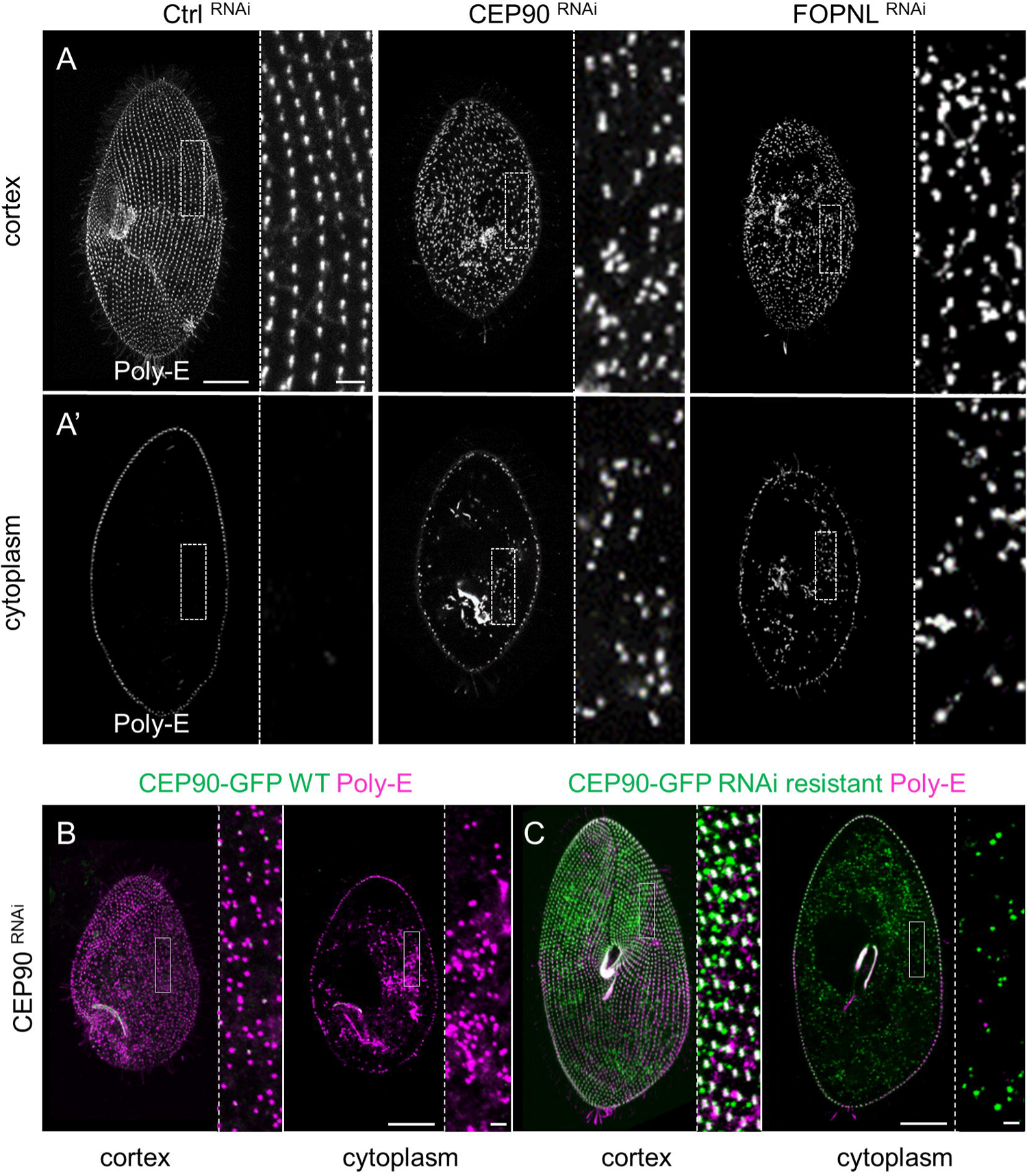
CEP90 depletion affects basal body docking. **A, A’**) Projection of confocal sections through the ventral surface (**A**) or at the intracytoplasmic level (**A’**), labeled with polyE antibodies (grey) in control (Ctrl) RNAi, *CEP90*^RNAi^ and *FOPNL*^RNAi^ *Paramecium*. In control-depleted cells, BB are organized in parallel longitudinal rows highlighted in the magnification (right panel) (**A**) with no BB observed in the cytoplasm (**A’**). Cells observed after 2 divisions upon CEP90 or FOPNL depletion appear smaller than control ones, with a severe disorganization of the BB pattern (**A**). Numerous internal BB are observed in the cytoplasm (**A’**). Scale bars= 20 µm and 2 µm (magnification). **B, C**) Specificity of *CEP90*^RNAi^: Transformants expressing either WT CEP90-GFP (**B**) or RNAi resistant CEP90-GFP (**C**) were observed after 2-3 divisions upon CEP90 depletion and analyzed for green emitted fluorescence and BB staining using poly-E antibodies. Scale bars= 20µm and 2 µm (magnification).The green fluorescence (green) of WT CEP90-GFP transformants was severely reduced along with the expected disorganisation of BB pattern at the cell surface (**B**). In contrast, the expression of RNAi-resistant CEP90-GFP rescued endogenous CEP90 depletion and restored the cortical organization of BB (**C**).

To analyze the effects of the depletion on the BB, inactivated cells were labeled for BB using poly-E antibodies. Whereas, control depleted cells (Ctrl^RNAi^) display BB at the cell surface organized in longitudinal rows (Figure 2A, cortex), Cep90-depleted cells show misaligned BB at the cortical surface (Figure 2A, CEP90^RNAi^). Confocal microscopy demonstrated that numerous BB were found in the cytoplasm after CEP90 depletion, suggesting BB anchoring defects (Figure 2A’, CEP90^RNAi^), while none were observed in control-depleted cells (Figure 2A’, Ctrl^RNAi^). The misaligned BB at the cell surface and cytoplasmic BB observed after CEP90 depletion was reminiscent of the phenotypes observed after depletion of either FOPNL or OFD1 (Figure 2 A, A’, FOPNL^RNAi^)(Aubusson-Fleury *et al*, 2012; Bengueddach *et al*, 2017). To ascertain the specificity of this phenotype, RNAi resistant CEP90a-GFP transformants were inactivated by CEP90 RNAi. GFP staining of RNAi resistant CEP90a-GFP transformants decorated BB as revealed by poly-E antibodies as CEP90a-GFP expressing paramecia (Figure 2C, cortex). In addition, cytoplasmic aggregates were observed, most probably due to the protein overexpression (Figure 2C, cytoplasm). Importantly, RNAi resistant CEP90a-GFP expression in paramecia inactivated for endogenous CEP90 rescued the BB organization at the cell surface, which displayed longitudinal rows of BB (Figure 2C). Finally, RNAi resistant CEP90a-GFP also rescued the dividing time, the survival and the swimming velocities to the level of control paramecia (Figure S3B, C). Altogether, these results suggest that CEP90 is involved in BB anchoring process in *Paramecium,* in agreement with the underlying ciliopathy phenotype observed in human.

Next, we used electron microscopy to analyze the ultrastructure of the undocked BB and better characterize the BB anchoring defect upon CEP90 depletion. Control-depleted cells show BB anchored at the cell surface and displaying the characteristic three plates of the TZ (Figure 3A, arrowheads). In contrast, numerous BB were lying in the cytoplasm in CEP90-depleted cells. All of them display defective distal ends with vestigial structures resembling the intermediate (blue arrowhead) and axosomal (magenta arrowhead) plates, while the terminal plate (green arrowhead) was mostly absent (Figure 3B-E). Occasionally, microtubule extensions were observed on the uncapped side of the BB (Figure 3E, white arrow).

**Figure 3:**
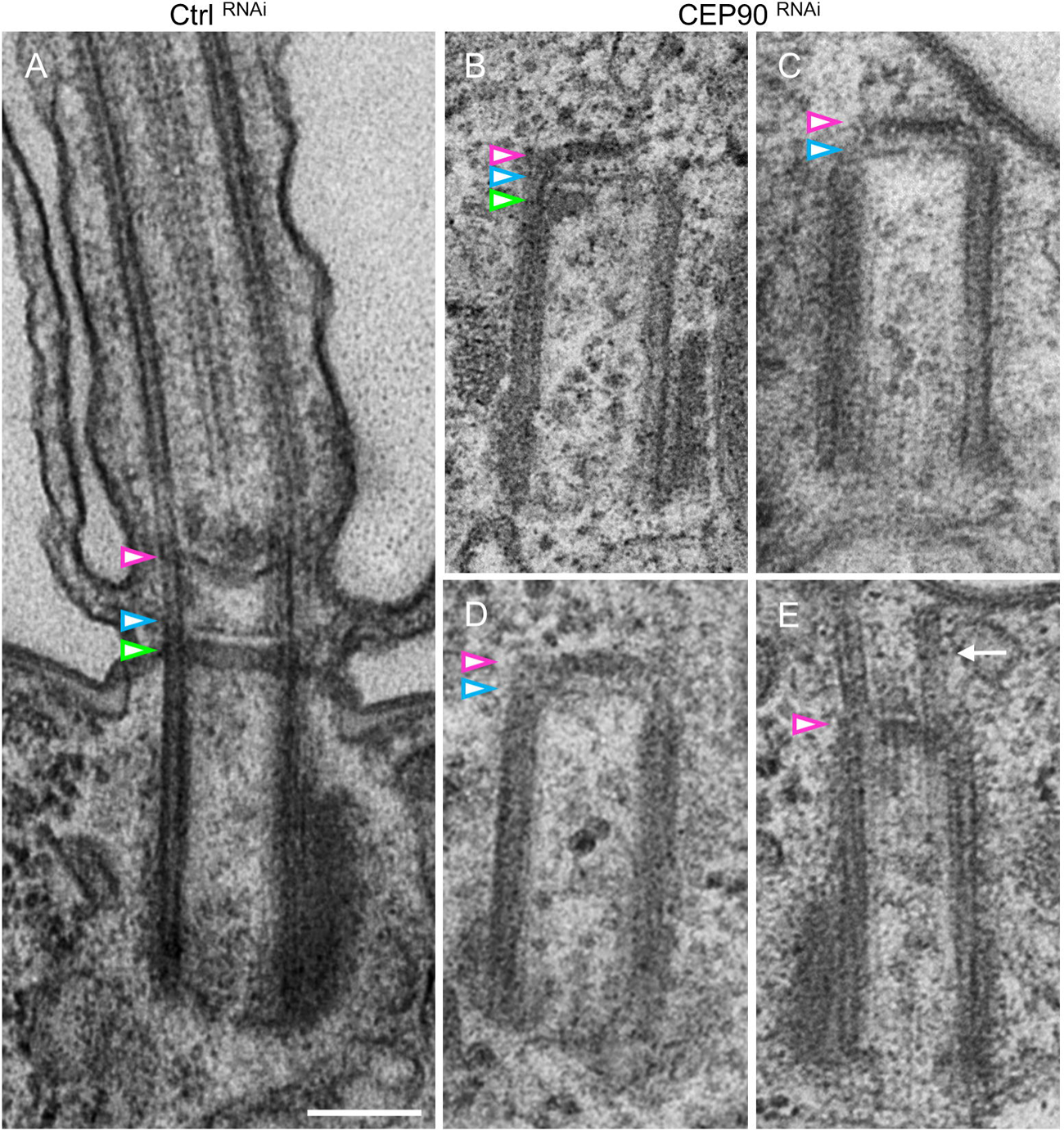
CEP90 depletion affects basal body’s distal end maturation. **A**) *Paramecium* showing one ciliated BB after control RNAi depletion (Ctrl). The TZ is characterized by three plates indicated by colored arrowheads: terminal plate (green), intermediate plate (blue) and axosomal plate (magenta). **B-E**) Upon CEP90 depletion, undocked BB are detected close to the cell surface (**C, E**) or deep in the cytoplasm (**B, D**). Vestigial structures resembling the axosomal plate (magenta arrowhead) and intermediate plate (blue arrowhead) are detected on all internal BB (n=63). **E**) Example of rare BB displaying microtubule extensions on the free, uncapped side of the BB (white arrow). Scale bar= 200nm.

Since, in mammalian cells, CEP90 was reported to affect centrosome duplication (Kodani *et al*, 2015), we investigated whether CEP90 depletion could also affect BB duplication in *Paramecium*. To do so, CEP90-depleted cells undergoing their first division and daughter cells resulting from this division were stained for BB. Intriguingly, we found that CEP90 depletion did not disturb BB duplication as demonstrated by the characteristic longitudinal groups of three or four BB issued from parental groups of two (Figure S4A). In the daughter cells, the presence of disorganized BB at the surface attests for an efficient CEP90^RNAi^ during this first duplication. We examine precisely the invariant field localized at the anterior part of these daughter cells, which is characterized by doublet BB units in the wild type cells. In this field, a defective BB duplication during division leads to singlet BB. As shown on Figure S4B, the invariant field encircled in yellow is mostly constituted of doublets BB in the absence of CEP90. Altogether, these results suggest that in *Paramecium* CEP90 depletion does not impair significantly the BB duplication process in contrast to its proposed role in mammalian cells (Kodani *et al*, 2015).

Taken together, these results indicate that CEP90 in *Paramecium* is required to build the distal ends of BB, which is necessary to dock the BB at the cell surface, as already observed for both FOPNL and OFD1.

### CEP90, FOPNL and OFD1 are required for basal body distal end assembly in Paramecium

In *Paramecium*, OFD1 and Centrin2 are required to assemble the BB distal end, as FOPNL (Bengueddach *et al*, 2017; Ruiz *et al*, 2005; Aubusson-Fleury *et al*, 2012). Centrin2 is recruited first and initiates the assembly of the three plates of the BB distal end. This step is followed by a co-recruitment of both FOPNL and OFD1 (Aubusson-Fleury *et al*, 2012; Bengueddach *et al*, 2017). Finally, Centrin3 is recruited to allow the BB tilting-up after duplication necessary for the its anchoring (Jerka-Dziadosz *et al*, 2013). To determine the involvement of CEP90 in distal end assembly, paramecia expressing GFP-tagged proteins (Centrin2, CEP90a, OFD1 or FOPNL) were inactivated by, either Centrin2, OFD1, FOPNL, CEP90 or Centrin3 RNAi. The fluorescence intensity in each transformed cells was quantified by immunostaining after two to three divisions under RNAi conditions (Figure 4 A,B). We found that the recruitment of CEP90a-GFP to BB, required the presence of FOPNL and OFD1 (Figure 4A,B). Reciprocally, CEP90 depletion prevented the recruitment of both OFD1-GFP (Figure 4A,B) and GFP-FOPNL (Figure S5A,B). These last results suggest an interdependent recruitment of OFD1, FOPNL and CEP90 to basal bodies. In addition, cells depleted for FOPNL, OFD1 or CEP90 maintained the recruitment of Centrin2 on newly formed BB (Figure 4A, B). Moreover, we demonstrated that Centrin2 depletion prevented the recruitment of CEP90 and OFD1 to the newly formed BB (Figure 4B, Figure S5C). By contrast, the depletion of Centrin3, which is not involved in the distal end assembly, did not affect the recruitment of Centrin2-GFP, CEP90a-GFP and OFD1-GFP (Figure 4 B, Figure S5D). Therefore, CEP90, FOPNL and OFD1 are co-recruited at the BB to assemble its distal end in *Paramecium*.

**Figure 4:**
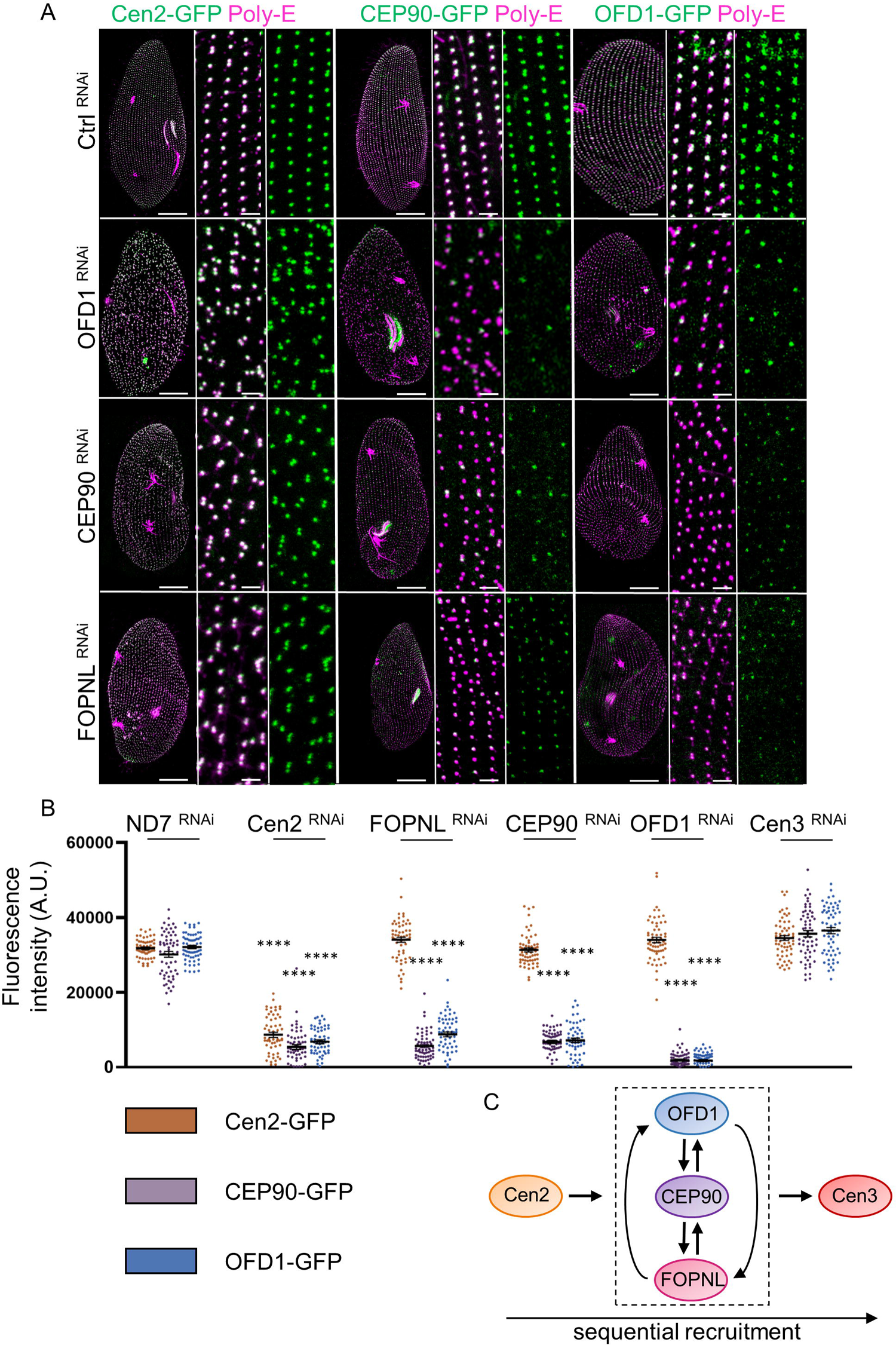
Sequential recruitment of basal body anchoring proteins in *Paramecium*. **A**) *Paramecium* transformants expressing either Cen2-GFP, CEP90-GFP or OFD1-GFP (green) and stained for BB (poly-E antibodies, magenta) after 2 or 3 divisions in control cells (Ctrl) and upon OFD1, CEP90 or FOPNL depletion. Control depleted paramecia displayed well anchored BB organized in parallel longitudinal rows, with the GFP emitted signal overlapping BB labeling. After OFD1, CEP90 and FOPNL depletion, BB pattern disorganization is observed. Scale bars= 20µm and 2µm (magnification). **B**) Quantification of the GFP fluorescence of CEP90-GFP (purple), Cen2-GFP (orange) and OFD1-GFP (blue) on newly formed BB in Control and upon Cen2, OFD1, FOPNL, CEP90, and Cen3 depletions. Dot plot showing the average intensity of GFP fluorescence (n= 80 cells, 3 independent replicates), AU: arbitrary units. Error bars show SEM. Statistical significance was assessed by one-way ANOVA followed by Tukey’s post hoc test ****p<0.0001.**C**) Schematic representation of the sequential recruitment of BB anchoring proteins leading to proper maturation of BB distal end. Cen2 arrives first at the BB followed by the interdependent recruitment of FOPNL, CEP90 and OFD1. This functional ternary complex is required to allow Cen3 recruitment and BB docking.

Altogether, these results show i) the similar localization of CEP90, FOPNL and OFD1 at BB distal extremity observed by U-ExM; ii) the interdependent recruitment of these 3 proteins; iii) their shared function in BB distal end assembly. This led us to propose that CEP90, FOPNL and OFD1 could form a functional ternary complex involved in BB distal end maturation and docking.

### Mammalian CEP90, OFD1 localize at the external surface of the distal end of mother and daughter centrioles as well as procentrioles

To investigate the functional conservation of CEP90 throughout the evolution, we analyzed CEP90 function in mammalian cells. In RPE1 cells, CEP90 localizes at centrioles and centriolar satellites, as previously shown by Kodani *et al*, (2015) reminiscent of FOPNL and OFD1 (Figure 5A, B and Figure S6A) (Sedjaï *et al*, 2010; Lopes *et al*, 2011). To further understand the organization of CEP90 in mammalian cells, we used U-ExM to resolve its localization at nanoscale resolution both in the osteosarcoma U2OS and RPE1 cell lines. We observed that, in cycling cells, the endogenous CEP90 or OFD1 longitudinal fluorescence signal is restricted to the centriolar distal end as compared to the tubulin signal, which depicts the whole centriolar length (Figure 5C, D, F and G; Figure S6B-C). The precise measurement of both CEP90 and OFD1 staining revealed that these two proteins localize on the two parental centrioles at about 375 nm from their proximal centriolar extremity (Figure 5C, F, L, M), and 50 nm from their distal end of the centrioles. From top-viewed centrioles, we observed a staining organized in a 9-fold symmetry close to the external surface of the centriolar microtubule wall (Figure 5 E, H). Since OFD1 and CEP90 antibodies were raised in rabbits, double labelling experiments could not be performed. Therefore, the measurements of both CEP90 and OFD1 staining relative to the tubulin maximal intensity signal show that CEP90 is slightly closer to the microtubule wall than OFD1 (average distance between CEP90 and tubulin about 30 nm, while OFD1 is about 40 nm) (Figure 5 L-N). Unfortunately, we have not been able to localize FOPNL using U-ExM, since our antibodies were not working under these conditions. Importantly, the analysis of both CEP90 and OFD1 staining during centriole duplication showed that both proteins were recruited on procentrioles (Figure 5C, D, F, G). Similar results were obtained in RPE1 cells (Figure S6D-F). In addition, ciliated RPE1 BB display a CEP90 staining similar to the unciliated ones (Figure S6G). Altogether, these results obtained by U-ExM demonstrate that the nanometric localization of both CEP90 and OFD1 in mammalian cells is similar to the one observed in *Paramecium*. The conserved localization between two evolutionary distant organisms might suggest a conserved function. We also investigated the localization of MNR, a direct interactor of OFD1 and FOPNL in mammalian cells (Chevrier *et al*, 2016). As expected, we found that MNR localizes at the distal end of the parental centrioles on the external surface of the microtubule wall in a 9-fold symmetry similarly as OFD1 and CEP90 (Figure 5I-K). This new result is in agreement with the relative position between OFD1, CEP90 and NMR (Kumar *et al*, 2021) but precisely unveils, for the first time, their position relative to the microtubule wall. Quantifications demonstrate that MNR localized in close proximity to the external part of the microtubule wall, next to CEP90 and OFD1 being slightly more external (Figure 5L-N). Observations in duplicating cells show the recruitment of MNR on early born procentrioles as observed for CEP90 and OFD1 (Figure S6B, C).

**Figure 5:**
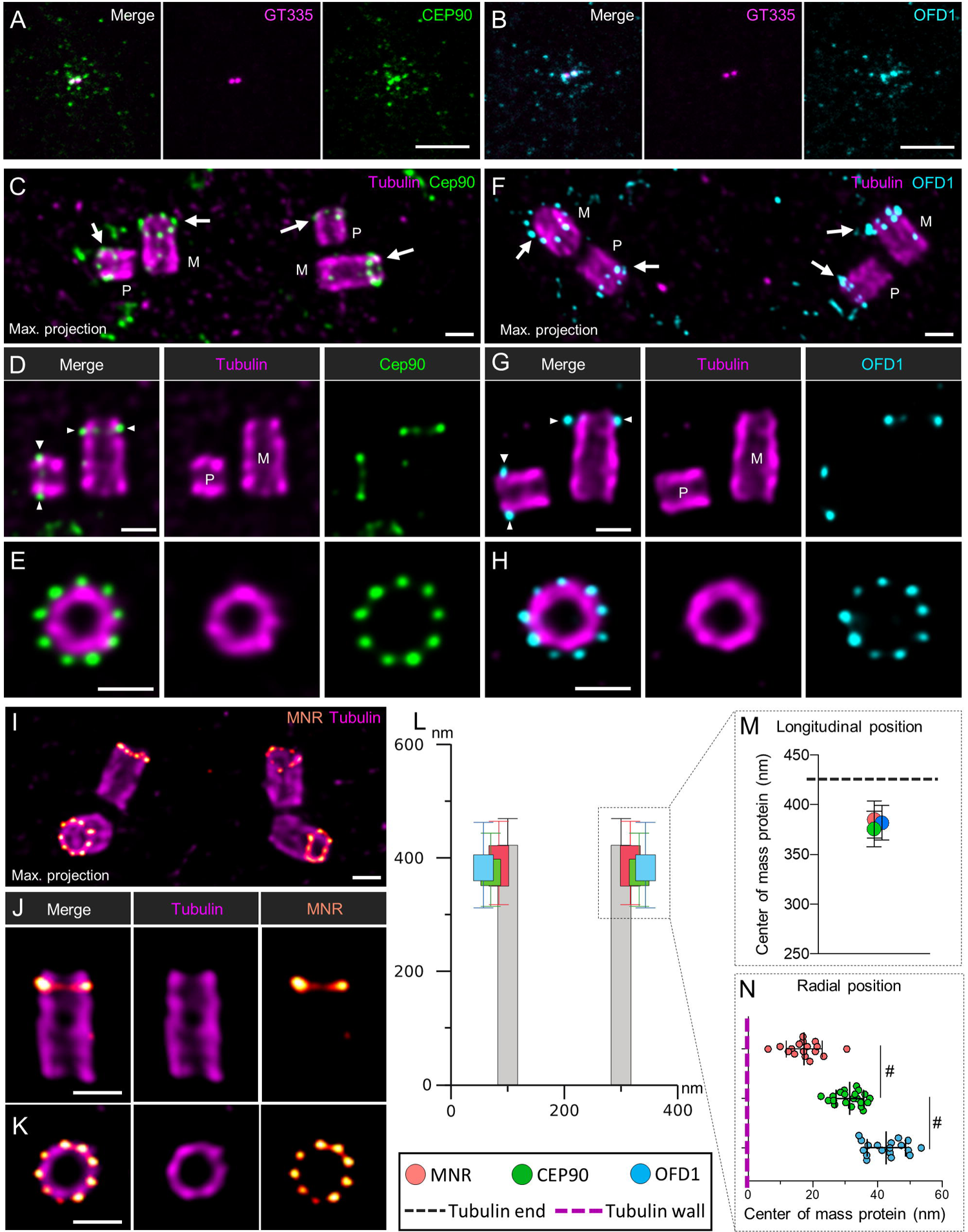
Centrosomal localization of CEP90 and OFD1 in mammalian cells. **A, B**) RPE1 cells were stained with anti-GT335 antibody directed against tubulin polyglutamylation (centrioles magenta) and anti-CEP90 (green) (**A**) or anti-OFD1 (cyan) (**B**). Both CEP90 and OFD1 decorated the centrosome and the centriolar satellites. Scale bar= 5µm. **C-J**) U2OS cells were expanded using the U-ExM technique and stained for tubulin (magenta) and CEP90 (green, **C-E**), OFD1 (cyan, **F-H**) or MNR (orange/red, **I-J**). Maximum intensity projections show the organization of CEP90 (**C**), OFD1 (**F**) and MNR (**I**) at the level of the centrosome in duplicating cells (M: Mature centriole; P: Procentriole). Single plan images shows that CEP90 (**D**), OFD1 (**G**) and MNR (**J**) localize slightly underneath the distal end of both the mother (M) as well as the procentrioles (P). Finally, images of top viewed oriented centrioles show 9-fold symmetry organization of CEP90 (**E**), OFD1 (**H**) and MNR (**K**) at the distal end of the centriole. Scale bars= 200nm. **L, M**) Distance between the proximal part of centriole (tubulin) and the fluorescence center of mass of CEP90 (green), OFD1 (cyan), or MNR (orange) (longitudinal position). N=23, 25, 49 centrioles for CEP90, OFD1 and MNR respectively from 3 independent experiments. Average +/- SD: CEP90= 375.3 +/- 17.8 nm, OFD1 = 381.4 +/- 17.4 nm, MNR = 385.1 +/- 18.7. **L, N**) Localization of CEP90, OFD1 and MNR with respect to the microtubule wall (radial position). Note that MNR localized the closest at the external surface of the microtubule wall, while OFD1 localizes more externally than CEP90. N=21, 18, 17 centrioles for CEP90, OFD1 and MNR respectively from 3 independent experiments. Average +/- SD: MNR = 17.4.1 +/- 5.5; CEP90= 31.4 +/- 4.3 nm, OFD1 = 42.7 +/- 5.9 nm. Statistical significance was assessed by Kruskal-Wallis test followed by Dunn’s post hoc test, CEP90 vs MNR p=0.06 (#), CEP90 vs OFD1 p=0.07 (#).

### The co-recruitment of CEP90, FOPNL, OFD1 is required for the formation of distal appendages

To investigate the hierarchy of recruitment of these three proteins to the centriole distal end in RPE1 cells. We first inactivated either *CEP90* or *FOPNL* gene using 2 sets of siRNA for each gene and found a significant reduction in CEP90 (80%) and FOPNL (80%) levels respectively at the centrosome (Figure 6A, B; Figure S7A) demonstrating the efficiency of the siRNA, as previously reported (Kodani *et al*, 2015; Sedjaï *et al*, 2010). Secondly, we analyzed the recruitment of CEP90, FOPNL and OFD1 to the centrioles upon depletion of CEP90 or FOPNL. We demonstrated that the recruitment of endogenous FOPNL and OFD1 at centrosomes was decreased of 65% upon CEP90 depletion (Figure 6B). Similarly, the depletion of FOPNL decreases the localization of both CEP90 and OFD1 to centrioles of about 70% and 60% respectively (Figure 6A, B). As a control of this experiment, we depleted either MNR or OFD1 by siRNA. In agreement with previous published data, depletion of either MNR or OFD1 in RPE1 cells prevents the recruitment of MNR, FOPNL and OFD1 (Chevrier *et al*, 2016) (Figure S7A’, B). In addition, both MNR and OFD1 depletion prevents also the recruitment of CEP90 (Figure S7A’, B) (Kumar *et al*, 2021). These results suggest that the interdependent recruitment of CEP90, FOPNL, OFD1 to centrioles and BB is conserved in *Paramecium* and mammalian cells.

**Figure 6:**
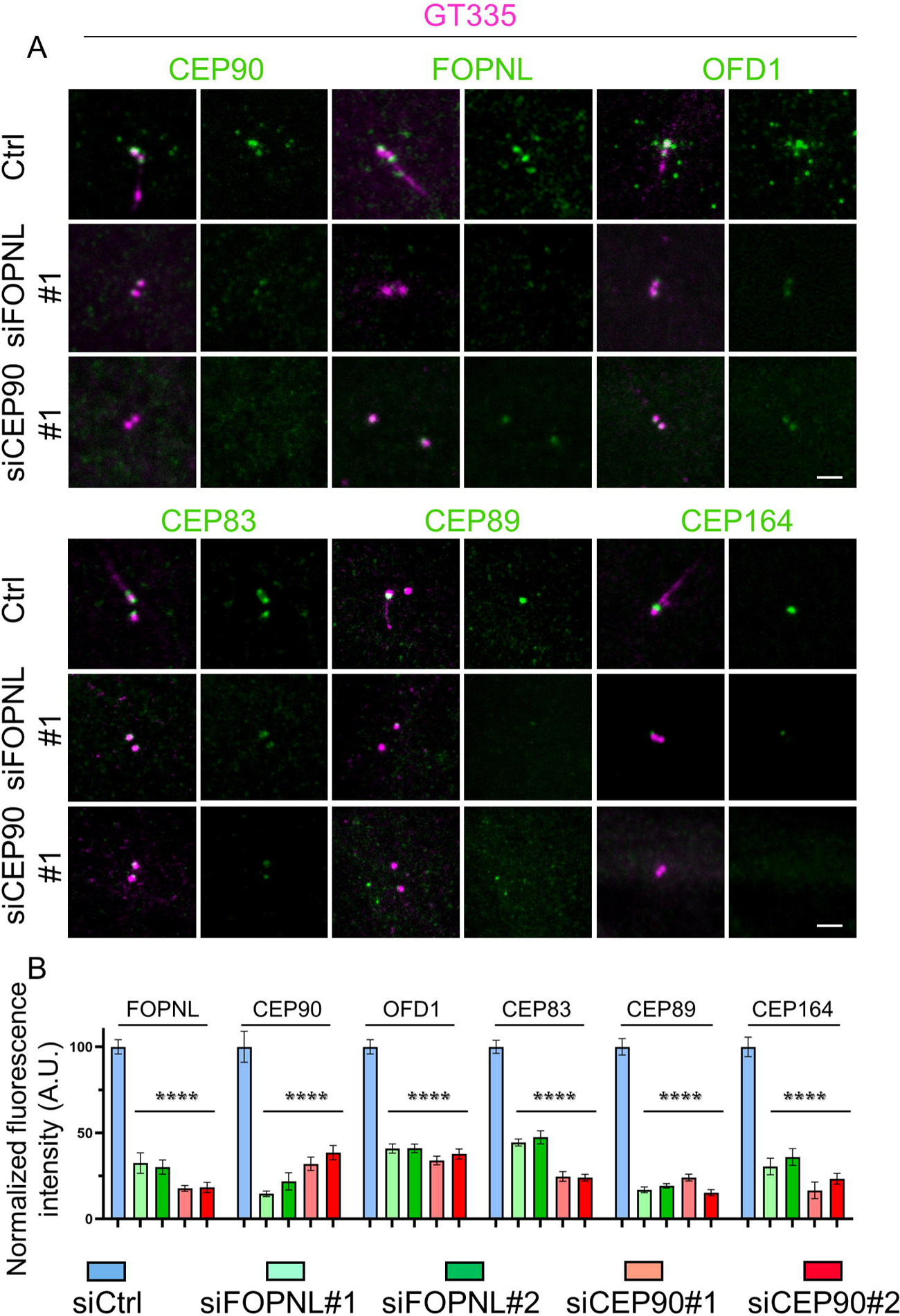
FOPNL and CEP90 regulate distal appendage proteins recruitment in mammalian cells. **A**) Serum starved RPE1 cells treated with control (ctrl), FOPNL or CEP90 siRNA were fixed and stained for GT335 (magenta) with CEP90, FOPNL, OFD1 or distal appendage proteins (CEP83, CEP89, and CEP164). Scale bar= 5µm. **B**) Quantification of the fluorescence intensity of FOPNL, CEP90, OFD1, CEP83, CEP89 and CEP164 at the centrosome. AU: arbitrary units. All data are normalized base on control siRNA and are presented as average ± SEM. ****p< 0.001 (one-way ANOVA followed by Tukey’s post hoc test), n ≥ 60 centrosomes from 2/3 independent experiments.

To gain further insights into the role of CEP90, OFD1 and FOPNL in ciliogenesis, we depleted either CEP90 or FOPNL proteins in serum-starved RPE1 cells. As expected from previous results (Kim *et al*, 2012; Sedjaï *et al*, 2010; Chevrier *et al*, 2016), we observed a significant decrease in cilia formation (about 80% compared to control siRNA) (Figure S8 A, A’, B) in both CEP90 and FOPNL siRNA treated cells.

Since we demonstrated, in *Paramecium,* that CEP90, FOPNL and OFD1 are involved in BB docking by controlling the assembly of the BB distal end, we investigated the formation of distal appendages after depletion of FOPNL in RPE1 to better understand their contribution in this process. Distal appendage formation requires both the presence of specific proteins able to recruit them and the removal of daughter specific proteins (Wang *et al*, 2018a). First, we analyzed if the depletion of CEP90 and FOPNL may affect the behavior of one daughter centriolar protein, Centrobin. As shown in Figure S8C, we did not observe any difference in the localization of Centrobin in control and CEP90 or FOPNL depleted cells. Secondly, we investigated the localization of the most proximal distal appendage protein CEP83, together with CEP89 and CEP164. Consistently, we found that the depletion of either CEP90 or FOPNL decreased the localization at the mother centriole of these distal appendage proteins of about 66% (Figure 6A, B). Altogether, these results suggest that the depletion of either CEP90 or FOPNL prevents the formation of the distal appendages, allowing the understanding of the defective ciliogenesis process previously observed (Wheway *et al*, 2015; Shen *et al*, 2020; Sedjaï *et al*, 2010; Xu *et al*, 2021). This suggest that FOPNL is a novel and important component for distal appendage assembly. As expected, depletion of OFD1 prevents distal appendage formation as observed by the poor recruitment of CEP83 and CEP164 as previously shown (Figure S7B) (Singla *et al*, 2010). Similar results were obtained for MNR depletion (Figure S7B)(Kumar *et al*, 2021). Consequently, BB docking cannot occur, leading to defective ciliogenesis.

### Overexpressed CEP90 recruits endogenous CEP83 to microtubules in cells overexpressing MNR

Our results obtained in mammalian cells showed that i) the complex composed of FOPNL, OFD1 and CEP90 is required for distal appendage formation; ii) The nanometric localization of of OFD1 and CEP90 is compatible with the localization of CEP83 along the BB proximo-distal axis (Bowler *et al*, 2019); iii) CEP90, OFD1 and MNR are recruited on early duplicating centrioles. Altogether, these results led us to propose the hypothesis that the FOPNL, OFD1, CEP90 complex will specify, from the procentriole stage, the future position of distal appendages which will be recruited on the daughter centriole during its maturation into a mother centriole. To test this hypothesis, we first analyzed the localization of CEP83 in U2OS cell lines by U-ExM. As expected, CEP83 is organized in a 9-fold symmetry at a similar position along the centriolar length as CEP90 and OFD1 (Figure 7A, B). Quantification of the average diameters of the rings revealed that CEP83 localized slightly more externally than OFD1 and CEP90. These results further support the hypothesis that FOPNL, OFD1 and CEP90 recruit distal appendage proteins to the centriolar wall. In order to decipher which protein of the ternary complex recruits the most proximal distal appendage protein, CEP83, we decided to take advantage of a displacement assay, in which overexpressed MNR-GFP localizes at microtubules and is used as a bait to displace other interacting proteins (Chevrier *et al*, 2016; Kumar *et al*, 2021). Therefore, MNR-GFP was overexpressed in U2OS alone or in combination with either myc-FOPNL, mcherry-OFD1 or CEP90. In each transfection condition, the putative displacement of endogenous OFD1, CEP90, MNR and CEP83 was first analyzed. As expected, expressed MNR-GFP decorated microtubules in U2OS cells (Figure 7F). Unfortunately, we could not displace any endogenous proteins with MNR as bait (Figure S9A). Reasoning that the amount of these proteins might be limited, we only looked at displacement of overexpressed protein by MNR-GFP. We demonstrated that MNR-GFP acts as a scaffold to recruit independently overexpressed OFD1, FOPNL and CEP90 (Figure 7G-L, Figure S9C, E, G). Importantly, endogenous OFD1 and CEP83 could be displaced on microtubules of the vast majority of the cell co-expressing CEP90 and MNR-GFP (Figure 7H, K). In contrast, only 10% of cells co-transfected with MNR-GFP and OFD1-mcherry were recruiting endogenous CEP90 and CEP83 (Figure 7I, L) to microtubules.

**Figure 7:**
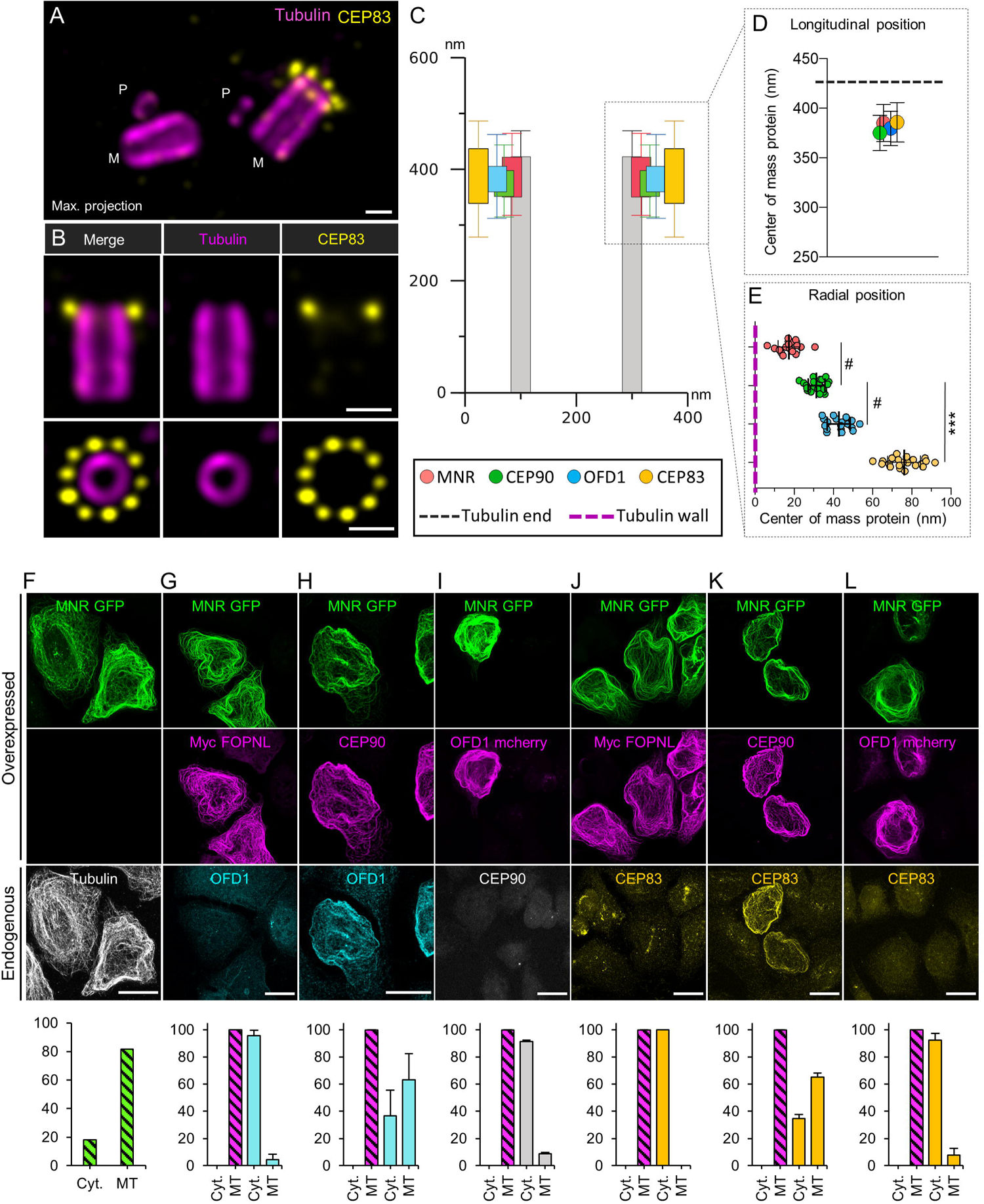
Recruitment of endogenous CEP83 on microtubules by overexpressed MNR-GFP and CEP90. **A-B**) U2OS cells were expanded using the U-ExM technique and stained for tubulin (magenta) and CEP83 (yellow). Maximum intensity projection shows the organization of CEP83, which is only present at the distal end of one of the two mature centriole and not at the level of the procentriole (M: Mature centriole; P: Procentriole) (**A**). Single plan images show that CEP83 localizes slightly underneath the distal end of both the mature centriole and is organized in 9-fold symmetry (**B**). Scale bars= 200nm. **C, D**) Distance between the proximal part of centriole (tubulin) and the fluorescence center of mass of CEP90 (green), OFD1 (cyan) MNR (orange) or CEP83 (yellow) (longitudinal position). N=23, 25, 49, 28 centrioles for CEP90, OFD1, MNR and CEP83 respectively from 3 independent experiments. Average +/- SD: CEP90= 375.3 +/- 17.8 nm, OFD1 = 381.4 +/- 17.4 nm, MNR = 385.1 +/- 18.7 nm, CEP83= 386.7 +/- 19.9 nm. **C, E**) Localization of CEP90, OFD1, MNR and CEP83 with respect to the microtubule wall (radial position). N=21, 18, 17, 21 centrioles for CEP90, OFD1, MNR and CEP83 respectively from 3 independent experiments. Average +/- SD: MNR = 17.4.1 +/- 5.5; CEP90= 31.4 +/- 4.3 nm, OFD1 = 42.7 +/- 5.9 nm, CEP83= 76.0 +/- 8.3 nm. Statistical significance was assessed by Kruskal-Wallis test followed by Dunn’s post hoc test, CEP90 vs MNR p=0.06 (#), CEP90 vs OFD1 p=0.07 (#), CEP90 vs CEP83 p<0.0001. **F-L**) U2OS cells transfected with MNR-GFP alone (**F**), MNR-GFP and Myc-FOPNL (**G, J**); MNR-GFP and CEP90 (**H, K**); and MNR-GFP and OFD1-mCherry (**I, L**). Transfected cells were stained with the following antibodies to visualize endogeneous proteins: anti-tubulin (**F**), OFD1 (**G, H**), CEP90 (**I**) and CEP83 (**J-L**). Scale bar= 20 µm. Cells co-expressed with either GFP-MNR/Myc; GFP-MNR CEP90; GFP-MNR/mCherry were analyzed for the localization of the endogenous OFD1, CEP90 and CEP83 proteins. The percentage of co-overexpressed protein localized to microtubules corresponds to 100% (striated bar) and the percentage of the endogenous proteins localized to cytosol (Cyt.) or microtubules (MT) for each condition. Blue bars correspond to endogenous OFD1; grey bars to CEP90 and yellow bars to CEP83. Averages and SDs are as follows: (**F**) Cyt.: 18.3%; MT: 81.7%, (**G**) Cyt.: 95.6%±4.1 MT: 4.4%±4.1, (**H**) Cyt.: 36,7% ± 19; MT: 63.3 %± 19, (**I**) Cyt.: 91.3%±1.2; MT: 8.7%±1.2, (**J**) Cyt.: 100%± 0; MT:0% ± 0, (**K**) Cyt.: 34.7%± 2.9; MT: 65.3± 2.9, (**L**) Cyt.: 92.3%± 4.9; MT: 7.7%± 4.9. 100<N <150 cells per condition from three independent experiments.

Altogether, these results suggest that MNR interacts, as previously shown, with OFD1 and FOPNL but also with CEP90. In addition, we discovered that CEP90 recruits CEP83, unveiling that the distal appendage proteins binding to the centriole occurs through CEP90.

## Discussion

In this study, using two complementary models, *Paramecium* and mammalian cells, we characterized the function of the FOPNL/OFD1/CEP90 complex in the BB anchoring process. We demonstrate that these proteins, together with MNR protein in mammalian cells, localize subdistally in a 9-fold symmetry at the external surface of the microtubule wall and determine the position and assembly of distal appendages.

Indeed, the search of FOPNL proximity partners in mammalian cells led us to identify CEP90. As expected from previous results (Chevrier *et al*, 2016; Kumar *et al*, 2021), we identified in our BioID mass spectrometry MNR/OFIP in proximity of FOPNL. Surprisingly, OFD1 is not significantly found despite the presence of biotinylated peptides. In addition, several centriolar satellite proteins as PCM1, CEP350, Cep131 or Talpid3 were significantly detected in proximity to FOPNL in our mass spectrometry, consistent with previous work (Gheiratmand *et al*, 2019b). CEP90 was previously identified as a centrosomal and centriolar satellite protein involved in spindle pole integrity in mitotic cells (Kim *et al*, 2012) or in centrosome duplication, by recruiting CEP63 to the centrosome (Kodani *et al*, 2015). CEP90 depletion in *Paramecium* could not recapitulate any defect in BB duplication, as we could always detect BB doublets in the invariant field of CEP90-depleted paramecium. We could speculate that, since *Paramecium* cells are devoid of centriolar satellites, the centriolar satellite pool of CEP90 in mammals is selectively involved in BB duplication, reinforcing previous results (Kodani *et al*, 2015). Another possible explanation could be that, despite numerous conserved actors (Jerka-Dziadosz *et al*, 2010; Gogendeau *et al*, 2011; Klotz *et al*, 2003; Ruiz *et al*, 2000; Dupuis-Williams *et al*, 2002), some proteins, such as CEP63, are not present in *Paramecium*, suggesting that the mechanism of BB duplication might differ from *Paramecium* to mammalian cells.

### CEP90, FOPNL and OFD1 are required for basal body anchoring

Patients with *CEP90* mutations display Joubert and Jeune syndrome, two ciliopathies. This result led Kim *et al* (2012) to postulate that CEP90 function in ciliogenesis may occur through centriolar satellites since their disruption inhibits the ciliogenesis process. To determine the function of CEP90 during ciliogenesis, we took advantage of the *Paramecium* model, which is physiologically devoid of centriolar satellites, simplifying the readout of the underlying phenotype of CEP90 depletion. We found that CEP90 depletion in *Paramecium* prevents BB distal end formation, as previously shown for FOPNL and OFD1, resulting in unanchored BB remaining in the cytoplasm. Interestingly, the depletion of CEP90 prevents maturation of the distal end of BB at the same step as FOPNL or OFD1 (Aubusson-Fleury *et al*, 2012; Bengueddach *et al*, 2017). This suggests that CEP90, FOPNL and OFD1 interact functionally to form a ternary complex involved in BB distal end assembly and in BB docking process. Our results obtained both in *Paramecium* and mammalian cells provide several arguments that may support this hypothesis. First, co-immunoprecipitation experiments performed in mammalian cells show an interaction between FOPNL, OFD1 and CEP90. Second, expansion microscopy performed both in *Paramecium* and mammalian cells, demonstrate that the three proteins localize similarly, close and external to the microtubule wall in a 9-fold symmetry. Third, the depletion of one of these proteins in *Paramecium* or mammalian cells prevents the recruitment of the two others. Finally, overexpressed MNR in mammalian cells recruits FOPNL, OFD1 or CEP90 suggesting that these 4 proteins constitute a functional complex.

In mammalian cells, the initial step of BB anchoring requires an interaction between the distal appendages and a cellular membrane (Knödler *et al*, 2010; Westlake *et al*, 2011; Schmidt *et al*, 2012; Sillibourne *et al*, 2013; Tanos *et al*, 2013; Shakya & Westlake, 2021). Remarkably, we found that either CEP90 or FOPNL depletion prevents the recruitment of distal appendage proteins, suggesting that the assembly of distal appendages is strongly impaired in these depleted cells as previously shown for OFD1 (Singla *et al*, 2010). Similar results were recently reported in a study performed in mammalian cells showing the role of the MNR, OFD1 and CEP90 complex in distal appendage assembly (Kumar *et al*, 2021).

Altogether, these results indicate that CEP90, FOPNL and OFD1 are required to tether the BB at a membrane both in *Paramecium*, bearing motile cilia, and in mammalian cells. Interestingly, *Paramecium* and mammalian cells use two distinct BB anchoring pathways. In *Paramecium*, the BB anchors directly to the cell membrane, whereas in RPE1 cells, cytoplasmic vesicles are recruited to the distal appendages, fuse to form the ciliary vesicle, in which the axoneme elongates, suggesting a functional conservation of this molecular complex in these two evolutionary distant organisms. Intriguingly, these proteins are absent in the *drosophila* and *C. elegans*. A possible explanation might be that in these species, the mechanism of BB anchoring might differ due to the absence of centriolar distal appendage structures. Interestingly, in *Drosophila* for instance, the distal appendage proteins CEP89, FBF1 and CEP164 are not required for BB docking and TZ formation, despite their recruitment, in a 9-fold symmetry, during BB anchoring (B. Durand personal communication).

### CEP90, FOPNL and OFD1 determine the future position of distal appendages assembly

The examination of the localization of CEP90, FOPNL and OFD1 by U-ExM in *Paramecium* and mammalian cells showed that these proteins localize at the distal end in a 9-fold symmetry at the external surface of the microtubule wall. Combining U-ExM with EM in *Paramecium*, we could determine that CEP90 remains at the same position along the MT while TZ maturation occurs. As previously described, a structural lengthening of the TZ occurs in *Paramecium* during ciliary growth (Gogendeau *et al*, 2020). CEP90 localizes to all BB whatever their ciliation status at the most proximal part of the TZ as observed by immunogold staining in EM. This is in contrast with the TZ proteins belonging to either MKS or NPHP complexes or the regulators of TZ assembly CEP290 and RPGRIP1L, which are recruited only at the distal part of the TZ of ciliated *Paramecium* BB. In *Paramecium*, the transition fibers emerge from the microtubule doublets at the level of the intermediate plate (Dute & Kung, 1978), a position compatible with our U-ExM and immunogold staining.

In agreement with the results obtained in *Paramecium*, U-ExM in mammalian cells showed that both the mother and the daughter centrioles were decorated. The localization of CEP90, OFD1 and MNR by U-ExM is observed slightly underneath the end of the centriole and at the external surface of the microtubule wall. In agreement with the results reported by Kumar and colleagues (Kumar *et al,* 2021), MNR diameter is the smallest one. However, co-staining of MNR and tubulin antibodies, which precisely determine the localization of the microtubule wall by U-ExM shows that MNR localization is external of the microtubule wall.

The localization of the CEP90, OFD1 and MNR complex is compatible with the localization of CEP83 along the BB proximo-distal axis (Bowler *et al*, 2019). Indeed, U-ExM staining of CEP83 in U2OS cell line demonstrates that it was the case. Examination of centrioles during the duplication process in both U2OS and RPE1 cell lines revealed that CEP90 and OFD1 are recruited on the newly born procentriole at a subdistal position similar to the one observed on mature centriole. This result differs from the one observed in Kumar *et al*, (2021), which reports the recruitment of the MNR, CEP90, OFD1 complex on elongating procentrioles during G2 phase. This discrepancy might be due to the fact that these authors used 5-ethynyl-29-deoxyuridine (EdU) to distinguish the different phases of the cell cycle, while we used unsynchronized U2OS and RPE1 cells. Consistent with our findings, numerous distal end proteins such as C2CD3, CP110, Centrin2 have already been observed on newly born procentrioles (Thauvin-Robinet *et al*, 2014; Kleylein-Sohn *et al*, 2007; Paoletti *et al*, 1996). However, the precise localization of these distal end proteins shows differences, C2CD3 beeing localized internally of the centriolar microtubule wall, while CP110 is at the extreme distal end. Therefore, the localization of the complex in agreement with the position of the proximal distal appendage protein CEP83 together with the early recruitment on procentriole led us to propose that the presence of the complex on the microtubule wall will mark the future position on which distal appendages will assemble. Accordingly, we used a displacement assay based on the overexpression of the microtubule-binding protein MNR to demonstrate that the co-expression of MNR and CEP90 is sufficient to recruit the most proximal distal appendage protein CEP83. Based on these results, we propose a model where MNR binds to the distal end of the centriolar microtubule wall and subsequently recruits OFD1, FOPNL and CEP90, to that location. Finally, CEP90 recruits CEP83. Altogether, the localization of the complex to procentrioles and the recruitment of endogenous CEP83 to co-overexpressed CEP90 and MNR proteins demonstrate that the complex composed of MNR, OFD1, FOPNL and CEP90 will dictate the future position of distal appendages (Figure 8).

**Figure 8:**
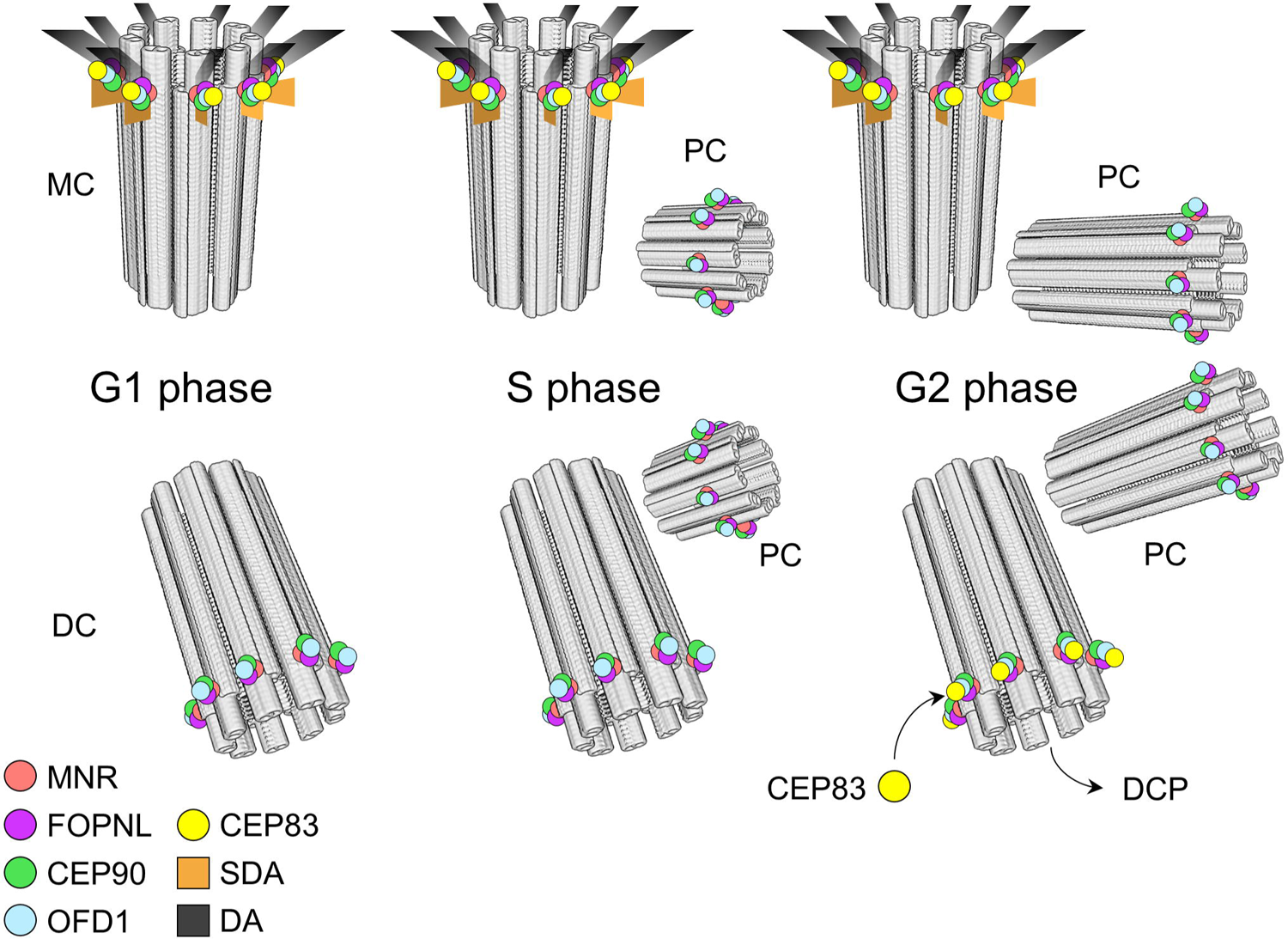
diagram of the function of the CEP90, FOPNL and OFD1 in distal appendages assembly. In G1, the complex composed of MNR, CEP90, FOPNL and OFD1 is localized to both mother (MC) and daughter (DC) centrioles. During centriole duplication (S phase), it is recruited on the two procentrioles (PC) at the distal extremity position, which is correlated with the future position of distal appendage protein CEP83. During G2/M the procentrioles elongate maintaining the complex distally. The ancient daughter centriole matures in a mother one, by losing DCP and assembling distal appendages, where the complex is localized. When one of the proteins of this complex (CEP90, FOPNL, and OFD1) is depleted, none of these proteins is recruited to the centriolar wall leading to defective distal end appendage assembly and defective ciliogenesis. We propose that this complex is required to dictate on the procentriole the future location of distal appendage assembly.

However, the reason for the localization of these proteins at the distal ends on both mother and daughter centriole is not yet clear. A possible explanation might come from the results of Wang et al, (2018), showing that daughter centriolar proteins (DCP) are observed on both mother and daughter centrioles after depletion of C2CD3 and Talpid3, which prevents distal appendages assembly, suggesting that DCP are removed from the mother centriole before distal appendages assembly. In contrast, the removal of OFD1 maintained the asymmetric localization of Centrobin on the daughter centrioles (Wang *et al*, 2018a). In agreement, the depletion of either CEP90 or FOPNL maintains the asymmetry of the two centrioles by maintaining Centrobin only at the daughter centriole. Therefore, we propose the following model: the ternary complex composed of CEP90, FOPNL and OFD1 is recruited soon after centrosome duplication on the two procentrioles, slightly underneath the distal end. POC1B (Chang *et al*, 2016) and POC5 (Azimzadeh *et al*, 2009) are then recruited to allow centriole elongation. Centrin2 (Ruiz *et al*, 2005; Aubusson-Fleury *et al*, 2012), OFD1 (Singla *et al*, 2010) and MNR (Kumar *et al*, 2021) have been shown to participate in the regulation of microtubule wall length, since their absence lead to elongated centrioles. Maturation of the daughter centriole in a mother one is carried out by first removing daughter centriole proteins and secondly by the recruitment of distal appendage proteins (Wang *et al*, 2018; Bowler *et al*, 2019; Viol *et al*, 2020) during G2/M (Figure 8). We propose that a similar scenario might also been conserved in *Paramecium* since the daughter centriole protein CEP120 is conserved. Intriguingly, according to our previous experiments, MNR is not functionally conserved in *Paramecium*. Therefore, how the complex FOPNL, OFD1 and CEP90 is tethered to the centriole wall is not clear. In mammalian cells, a functional interaction between C2CD3 and OFD1 has been reported (Ye *et al*, 2014; Wang *et al*, 2018b). An orthologue of C2CD3 has been found in *Paramecium* (Zhang & Aravind, 2012). A possibility would be that OFD1 is tethered to the centriolar wall by C2CD3 and will recruit the other member of the complex with CEP90 recruiting distal appendages proteins.

To conclude, the OFD1, CEP90 and FOPNL complex appears as the core evolutionary conserved complex necessary to allow distal appendage location and assembly both in primary cilia and motile cilia.

## Material and methods

### P. tetraurelia strains and cultures conditions

The wild type reference strain, stock d4-2 of *Paramecium tetraurelia*, was used in RNAi experiment. To facilitate the screening of transformants in microinjection experiments, the mutant *nd7-1* (Skouri & Cohen, 1997) which carries a mutation preventing trichocyst exocytosis was used. Cells were grown at 27°C in wheatgrass infusion (BHB) bacterized with Klebsiella pneumoniae and supplemented with 0.8 µg/mL of β-sitosterol according to standard procedures(Sonneborn, 1970).

### Human cell lines and culture condition

The human retinal pigment epithelium (RPE1), human embryonic kidney (HEK293), human bone osteosarcoma (U2OS) cell lines were grown in DMEM/F12 (RPE1), DMEM (HEK293, U2OS), respectively supplemented with 10% of Fetal bovine serum and Penicillin-streptomycin (1000 units/mL and 1000µg/mL respectively HyClone), Glutamax cultured in the presence of 5% CO2. Flp-In T-REx HEK293 cells were grown in DMEM supplemented with 10% FBS, GlutaMAX, zeocin (100 μg/mL), and blasticidin (3μg/mL). Flp-In T-REx HEK293 stable lines expressing myc-BirA* or myc-BirA*-FOPNL were maintained as above, with the addition of hygromycin (200 μg/mL) instead of zeocin.

### Antibodies

The following antibodies were used: the polyclonal (1/100) and monoclonal (gt335, 1/10000) PolyE antibody (Janke & Bulinski, 2011) stained cilia. The monoclonal anti-epiplasmin (V3E2)(Aubusson-Fleury *et al*, 2013). For mammalian cells, polyclonal rabbit anti-CEP90 (1/250 or 1/500-Proteintech 14413-1-AP ((Kodani *et al*, 2015)), anti-OFD1 (1/250 or 1/500, Sigma HPA031103 (Wang *et al*, 2013)), anti-CEP83 (1/250 or 1/500-Proteintech 26013-1AP, (Hossain *et al*, 2020)), anti-MNR/KIAA0753 (1/250 - Novusbio NBP1-90929) anti-Cep89 (1/800-Proteintech 24002-1AP, (Kanie *et al*, 2017)), Cep164 (1/1000-Proteintech 22227-1AP, (Bowler *et al*, 2019)), anti-Centrobin (1/500-mouse-Abcam-Ab70448 (Wang *et al*, 2018a) and monoclonal anti-FOPNL antibodies (1/200-(Sedjaï *et al*, 2010)) were used. Rat monoclonal anti-MNR were used (1/500; Chevrier *et al*, 2016) for Microtubule displacement assays while anti MNR (Novus Biological NBP1-90929) for expansion microscopy. Monobodies AA344 (1:250, scFv-S11B, Beta-tubulin) and AA345 (1:250, scFv-F2C, Alpha-tubulin; (Nizak *et al*, 2003) were used in expansion microscopy to stain for the microtubule wall. Goat anti-rabbit Alexa Fluor 488 F(ab’)2 fragment and goat anti mouse Alexa Fluor 568 IgG (H+L) were provided by Invitogen and used at 1/1000.

### BioID sample preparation

Since we searched for proteins at proximity of the centrosomal pool of FOPNL, we first enriched our sample in centrosomes using isolated centrosome-nucleus complexes (Maro & Bornens, 1980) from HEK293 FlpIn mycBirA*FOPNL and HEK293 FlpIn mycBirA* cells in presence or absence of Tetracycline. BioID purification was then performed according to (Roux *et al*, 2012). Briefly, the purified centrosome-nucleus complexes were then incubated 10 min in 50mM Tris pH 7.4, 300mM NaCl, 0.4% SDS, 5mM EGTA, 1mM DTT, Complete anti-protease cocktail (Roche) then 1% Triton-X-100 were added for 10 min. An equivalent volume of Tris pH 7.4 was added and the lysate was centrifuged à 4°C at 25 000g for 10min at 4°C. The supernatant was collected and incubated overnight with 70 µL of streptavidin beads with gentle agitation at 4°C. Streptavidin beads were washed with 50mM Tris pH7.4. After the last wash, the proteins were recovered from the beads using Laemmli sample buffer. Proteins were allowed to enter the SDS acrylamide gel for 1cm. Gels were stained using colloidal blue. The gel containing the proteins is then cut (gel plug). 3 independent replicates were done for myc BirA* cells and 2 independent replicates for mycBirA*FOPNL.

### Samples preparation prior to LC-MS/MS analysis

Gel plugs were discolored using a solution of ACN/NH_4_HCO_3_ 50mM (50/50) during 15 min with agitation. Plugs were reduced with a solution of 10 mM DL-Dithiothreitol (DTT, Sigma-Aldrich, Saint Louis, MO, USA) during 45 min at 56°C, then the plugs were alkylated using à 55mM solution of Iodoacetamide (IAA, Sigma-Aldrich, Saint Louis, MO, USA) during 45 min at room temperature. After a step of washing and dehydration, proteins in the plugs where digested overnight with Trypsin (10μg/ml) (Promega, Madison, WI, USA) at 37°C in a 25-mM NH_4_HCO_3_ buffer (0.2 µg trypsin in 20 µL). The resulting peptides were desalted using ZipTip μ-C18 Pipette Tips (Pierce Biotechnology, Rockford, IL, USA).

### LC-MS/MS acquisition

Samples were analyzed using an Orbitrap Fusion equipped with an easy spray ion source and coupled to a nano-LC Proxeon 1200 (Thermo Scientific, Waltham, MA, USA). Peptides were loaded with an online preconcentration method and separated by chromatography using a Pepmap-RSLC C18 column (0.75 x 750 mm, 2 μm, 100 Å) (Thermo Scientific), equilibrated at 50°C and operating at a flow rate of 300 nl/min. Peptides were eluted by a gradient of solvent A (H_2_O, 0.1 % FA) and solvent B (ACN/H_2_O 80/20, 0.1% FA), the column was first equilibrated 5 min with 95 % of A, then B was raised to 28 % in 105 min and to 40% in 15 min. Finally, the column was washed with 95% B during 20 min and re-equilibrated at 95% A during 10 min. Peptide masses were analyzed in the Orbitrap cell in full ion scan mode, at a resolution of 120,000, a mass range of *m/z* 350-1550 and an AGC target of 4.10^5^. MS/MS were performed in the top speed 3s mode. Peptides were selected for fragmentation by Higher-energy C-trap Dissociation (HCD) with a Normalized Collisional Energy of 27% and a dynamic exclusion of 60 seconds. Fragment masses were measured in an Ion trap in the rapid mode, with and an AGC target of 1.10^4^. Monocharged peptides and unassigned charge states were excluded from the MS/MS acquisition. The maximum ion accumulation times were set to 100 ms for MS and 35 ms for MS/MS acquisitions respectively.

### Data analysis LC-MS/MS

Label-free quantification was done on Progenesis QI for Proteomics (Waters, Milford, MA, USA) in Hi-3 mode for protein abundance calculation. MGF peak files from Progenesis were processed by Proteome Discoverer 2.2 with the Mascot search engine (Matrix Science, version 2.4). Identification of proteins was done with the Swissprot database <u>release 2018_</u>12 with the *Homo sapiens* taxonomy. A maximum of 2 missed cleavages was authorized. Precursor and fragment mass tolerances were set to respectively 7 ppm and 0.5 Da. The following post-translational modifications were included as variable: Acetyl (Protein N-term), Oxidation (M), Phosphorylation (STY), D-Biotin (K). The following post-translational modifications were included as fixed: Carbamidomethyl (C). Spectra were filtered using a 1% FDR using the percolator node. The mass spectrometry proteomics data have been deposited to the ProteomeXchange Consortium via the PRIDE (Perez-Riverol *et al*, 2019) partner repository with the dataset identifier PXD028740”. We would recommend you to also include this information in a much abridged form into the abstract itself, e.g. “Data are available via ProteomeXchange with identifier PXD028740.”

### CEP90 gene identification in Paramecium

By BLAST search in Parameciumdb (https://paramecium.i2bc.paris-saclay.fr/), we identified two *Paramecium* genes encoding proteins homologous to CEP90: *CEP90a* (PTET.51.1.G0320241) and CEP90b (PTET.51.1.G0320283) (Arnaiz *et al*, 2020). CEP 90a and b protein shares about 20,7% identity with human CEP90 protein. These two genes result from the last whole genome duplication, which occurred in *Paramecium*, and share 95,2% identity. RNAseq analysis as well as microarray show that *CEP90a* is 10x more expressed than *CEP90b*.

### Genes cloning

For gene silencing, the more divergent part of the gene was chosen to selectively inactivate either *CEP90a* or *CEP90b*, since the two paralogs of *CEP90* in *Paramecium* share a high percentage of identity. The inactivation of both CEP90a and CEP90b was performed using a L4440 vector containing the same *CEP90a* fragment (1472-1951 from ATG start codon) linked to a new fragment of *CEP90b* (139-618 from ATG start codon). Since the 139-618 *CEP90b* fragment is localized in a less divergent part, an OFF-Target effect has been identified on *CEP90a* with a hit of 3. Silencing vectors of FOPNL, OFD1, PtCen2 and PtCen3 have already been described previously (Aubusson-Fleury *et al*, 2012; Bengueddach *et al*, 2017; Ruiz *et al*, 2005). This vector allows the synthesis of double-stranded RNA corresponding to the cloned gene from two T7 promoters. The possible off-target effect of this sequence was analyzed using the RNAi off-target tool in the *P. tetraurelia* genome database (https://paramecium.i2bc.paris-saclay.fr/cgi/tool/rnai_off_target) that searches the genome for stretches of 23 identical nucleotides (the size of *Paramecium* siRNA). The only genomic sequence matching for *CEP90a* and *CEP90b* RNAi fragment were *CEP90a* or *b*, thus ruling out possible off-targeting.

For CEP90-GFP expression, *CEP90a* gene showing a 10x higher expression level than *CEP90b* gene during the vegetative state of *Paramecium*, *CEP90a* gene was chosen for expressing the GFP-tagged fusion protein. A synthetic *CEP90a* gene, resistant to *CEP90a* RNAi was designed by punctual nucleotide modifications (from base 1472 to 1951 from the ATG start codon). Both wild type and RNAi resistant constructions were inserted in PPXV-GFP plasmids. As a result, the GFP is fused in the 3’end of *CEP90a* and RNAi resistant *CEP90a* gene. The fusion proteins are expressed under the control of the calmodulin promoter.

Human cDNA of FOPNL (provided by O. Rosnet) was amplified by PCR using primers (Table S2) and cloned into pcDNA5 FRT/TO mycBirA* using the appropriate restriction sites. The plasmids MNR-GFP, OFD1-mcherry (Chevrier *et al*, 2016) as well as CEP90 (clone ID 6140514, Dharmacon horizondiscovery).

### Generation of stable and inducible HEK293 flpIn FOPNL for BioID

Flp-In T-REx HEK293 cells were cotransfected with pOG44 (Flp-recombinase expression vector) and either pcDNA5FRT/TO mycBirA*FOPNL or pcDNA5FRT/TO. Transfections were performed with Lipofectamine2000 according to the manufacturer’s instructions. After transfection, cells were selected with hygromycin (200µg/mL) and blasticidin (15µg/mL). HEK293 FlpIn mycBirA*FOPNL and HEK293 FlpIn mycBirA* cells were incubated for 24 h with 50 μM biotin (Sigma-Aldrich) in the presence or absence of 0.1 μg/ml tetracycline.

### Paramecium transformation

*Nd7-1* cells were transformed by microinjection into their macronucleus (Gilley *et al*, 1988). DNA containing a mixture of the plasmid of interest (*CEP90-GFP*, *CEP90RNAi resistant GFP*, *OFD1-GFP*, *GFP-FOPNL* at 5 μg/μl) with the DNA directing the expression of the *ND7* gene to complement the *Nd7* mutation. Microinjection was made under an inverted Nikon phase-contrast microscope, using a Narishige micromanipulation device and an Eppendorf air pressure microinjector. Transformants were screened for their ability to discharge their trichocysts and further analyzed for GFP. Clones expressing the fusion protein and showing a growth rate similar to untransformed paramecia were chosen for further analyses.

### Gene silencing in Paramecium by feeding

L4440 inactivation vectors were introduced in HT115 DE3 *Escherichia coli* strain to produce T7Pol-driven dsRNA as previously described (Galvani & Sperling, 2002). Paramecia were fed with these bacteria and refed daily with a fresh feeding medium at 27°C. Control cells were fed with HT115 bacteria carrying the L4440 vector containing the *ND7* gene (control). Phenotypes were analyzed after 48h of feeding. In these conditions, most of the mRNA of silenced genes are knocked-down therefore leading to the depletion of the proteins.

Effectiveness of RNAi was quantified by analyzing the decrease of BB fluorescence in transformed cell lines expressing the GFP-tagged transgene after silencing. In *Paramecium*, the phenotype of silenced cells is highly reproducible from one experiment to another and from one cell to another. In each experiment paramecia were analyzed in at least 2 or 3 independent experiments.

### Paramecia fixation for immunofluorescence

Fixation and immunofluorescence techniques were performed on cells in suspension. 50-100 cells were collected in the smallest volume possible and were permeabilized in 200 µl PHEM (Pipes 60mM, Hepes 25 mM, EGTA 10mM, MgCl2 2mM, adjusted to pH 6.9) with 1% Triton-X-100 (PHEM-Triton) for 30 secondes. Cells were fixed for 10min in 2% PHEM-PFA. Buffer was then aspirated and cells were rinsed 3 times for 10 min in PBS –tween 0.1%-3% BSA

### Confocal acquisition

Confocal acquisitions were made with a Leica SP8 equipped with a UV diode (line 405), and three laser diodes (lines 488, 552 and 635) for excitation and two PMT detectors. For U-ExM data, images were collected with an inverted confocal Microscope Leica DMI 6000 CS (for *Paramecium cells*) or an inverted Leica TCS SP8 microscope (for human cells) using a 63 × 1.4 NA oil objective with Lightening mode at max resolution, adaptive as ‘Strategy’ and water as ‘Mounting medium’ to generate deconvolved images. 3D stacks were acquired with 0.12 µm z-intervals and an x, y pixel size of 35 nm.

### Expansion microscopy-U-ExM

For analyzing CEP90 and OFD1 in human centrioles, U-ExM was performed as previously described (Steib *et al*, 2020). The following reagents were used in U-ExM experiments: formaldehyde (FA, 36.5–38%, F8775, SIGMA), acrylamide (AA, 40%, A4058, SIGMA), N,N’-methylenbisacrylamide (BIS, 2%, M1533, SIGMA), sodium acrylate (SA, 97–99%, 408220, SIGMA), ammonium persulfate (APS, 17874, ThermoFisher), tetramethylethylendiamine (TEMED, 17919, ThermoFisher), nuclease-free water (AM9937, Ambion-ThermoFisher) and poly-D-Lysine (A3890401, Gibco). Briefly, coverslips were incubated in 2% AA + 1.4% FA diluted in PBS for 5 hr at 37°C prior to gelation in monomer solution (19% sodium acrylate, 0.1% BIS, 10% acrylamide) supplemented with TEMED and APS (final concentration of 0.5%) for 1 hr at 37°C and denaturation for 1h30 min at 95°C, gels were stained for 3 hr at 37°C with primary antibodies: CEP90, OFD1, CEP83 and MNR were used with a combination of tubulin monobodies (see antibodies section) diluted in PBS-BSA 2%. After 3 PBS-tween 0.1% washes, secondary antibodies were incubated for 2h30 min at 37°C. Gels were then washed 3 times in PBS-tween 0.1% and incubated in water overnight to allow expansion.

For *Paramecium* BB expansion microscopy, the following modification has been done: purified cortex was incubated in 1%AA + 0.5%FA diluted in 1XPBS for 5h at 37°C. The GFP-tagged proteins were stained with Rabbit anti-GFP (1/400 Torrey Pines TP-401) and the combination of tubulin monobodies as above.

### Electron microscopy

For ultrastructural observations, cells were fixed in 1% (v/v) glutaraldehyde and 1% OsO4 (v/v) in 0.05 M cacodylate buffer, pH 7.4 for 30 min. After rinsing, cells were embedded in 2% agarose. Agarose blocks were then dehydrated in graded series of ethanol and propylene oxide and embedded in Epon812 (TAAB, Aldermaston, Berkshire, UK). For pre-embedding immunolocalization, the immunostaining process was carried out as described for immunofluorescence using gold-coupled instead of fluorochrome-coupled secondary antibodies (gold-labeled anti-rabbit IgG-GAR G10, Aurion) diluted 1/30 for 30 min. Cells were then treated as described above.

All ultrathin sections were contrasted with uranyl acetate and lead citrate. The sections were examined with a Jeol transmission electron microscope1400 (at 120kV).

### Paramecium swimming analysis

4 to 8 paramecia were transferred in 10 µl drops of conditioned BHB (bacterized BHB solution which has been depleted for bacteria after their growth and sterilized) for 15 min before being tracked for 10s every 0.3s. We used a Zeiss Steni 2000-C dissecting microscope with a Roper Coolsnap-CF camera and Metamorph software (Universal Imaging). Stacks were analyzed using the Manual tracking tool in ImageJ.

### FOPNL immunoprecipitations

Hela Kyoto TransgeneOmics cells expressing GFP-tagged mouse FOPNL were grown, washed in phosphate buffered saline (PBS) and lysed in 50 mM Tris–HCl pH 7.5, 150 mM NaCl, 1 mMEDTA, containing 1% NP-40 and 0.25% sodium deoxycholate plus a complete protease inhibitor cocktail (Roche) on ice. Clear lysates were obtained after centrifugation 15 minutes at 13000rpm at 4°C. To immunoprecipitate FOPNL, cell lysate was incubated with agarose beads conjugated with either mouse IgG or mouse anti-GFP antibodies (Roche) for 3h at 4°c under gentle agitation. Beads were washed 5 times with lysis buffer and finally resuspended in Laemli sample buffer. Proteins were separated by SDS-PAGE and transferred onto nitrocellulose. Blots were revealed with rabbit anti GFP, rabbit CEP90 and rabbit OFD1 antibodies.

### Mammalian cell fixation and immunofluorescence

RPE1 cells were grown on coverslips, washed in PBS, and fixed in methanol at −20°C for 6 min. Alternatively, after a PBS wash, cells were extracted in a PHEM buffer (45 mM Pipes, 45 mM Hepes, 10 mM EGTA,5 mM MgCl2 adjusted to pH 6.9, and 1 mM PMSF) containing 1% TritonX-100 and then fixed as described above. Cells were rinsed in PBS containing 0.1% Tween 20. Primary antibodies diluted in PBS containing 3%BSA were added for 1 h at room temperature. Three washes were performed in PBS-Tween, and the Alexa-labeled secondary antibodies were applied (FisherScientific). Cells were washed in ethanol and mounted in Citifluor (CityUniversity, London, England). Observations were done on a confocal microscope. All figures presented are two-dimensional projections of images collected at all relevant z-axes

### siRNA-mediated silencing in mammalian cells

30000 RPE1 cells were plated onto coverslips 24 hr prior transfection. For CEP90 depletion, two siRNA were used. One is a smart pool of Dharmacon (horizondiscovery) and the other one corresponding to nucleotides 1651-1675 and used in the Kodani’s paper (Kodani *et al*, 2015). Similarly, two siRNA were used for FOPNL. One is a smart pool of Dharmacon and the other one the published siRNA sequence (Sedjaï *et al*, 2010) as well as a smart pool of Dharmacon. Scrambled siRNA was used as a negative control. Cells were transfected with 20nM of siRNA using Lipofectamine RNAiMax (Invitrogen). Medium was changed after 48 hr post-transfection tio induce ciliation and cells were analyzed 72 hr post-transfection.

### Microtubule recruitment assay

60 to 70% confluent U2OS cells grown on coverslips were transfected in a six-well plate using lipofectamine 2000 following the manufacturer’s instructions, with 1µg of total DNA of the following combinations: MNR-GFP alone, OFD1 mCherry alone, CEP90 alone, myc-FOPNL alone, MNR-GFP with either myc-FOPNL, OFD1 mCherry or CEP90. Expression of the different fusion proteins was allowed for 24 hours. Cells were then fixed for 3 min in −20°C cold MeOH and washed once in PBS before staining with antibodies against OFD1, CEP90, myc, CEP83. The GFP and mcherry emitted fluorescence was observed directly. After three washes in PBS Tween 0.1%, coverslips were mounted using glycerol-mounting medium and DABCO 1, 4-diazabicyclo (2.2.2) octane (Abcam, ab188804).

### Image analysis

For immunofluorescence, image stacks were processed with ImageJ and Photoshop. Confocal acquisition parameters were kept identical (same laser intensity and same gain and same resolution) for quantification of BB fluorescence intensity in control and RNAi treated cells. Average pixel fluorescence intensity of the different antibodies was then measured within a defined and constant circular area of the size of the BB (*Paramecium*) or centrosome (RPE1 cells), using the measurement tools of ImageJ.

For U-ExM data acquisition of Paramecium cortex, confocal images were acquired with a LEICA TCS SP8x (DMi 6000) equipped with a 63x Apochromat oil-immersion objective (numerical aperture (NA): 1.4), U.V diode laser 405nm and a white light laser (WLL) AOTF coupled. Excitation at 490nm and 571nm of Alexa 488 and Alexa 568, respectively. Images were recorded with two hybrid detectors. Acquistions were made at 512×512 pixels and 400Hz speed and zoom of 8.4. Alexa 488 was acquired with a frame accumulation of 3. Sample sampling was 43nm in XY and for 3D imaging, stacks were acquired with a z-step of 130nm. The whole system was driven by LAS X software version 3.5.6. Image manipulation were done with ImageJ (Wayne Rasband, National Institutes of Health). Images were deconvolved employing the Huygens Essential package (Scientific Volume Imaging, Hilversum, The Netherlands), with sampling of 43nm in XY and 130nm in Z and the following parameters: algorithm: CMLE; signal/noise ratio: 20; quality threshold: 0.01%; maximum iterations: 40; background calculation: in/near objects.

For U-ExM analysis, length coverage and diameter quantification was performed as previously published in (Steib *et al*, 2020b).

To generate the panels in Figure 1F, 1G, 5L, 5M, 5N, 7C, 7D, and S2F, two homemade plugins for ImageJ were used. The first one helps the user to easily pick the extremities of a centriolar protein signal. For each protein of each image, the user picks the first and the last point of the signal along the centriole length. This generates an entry in a table for each centriole describing where the proteins start and end. The results are saved in a .csv file that can be directly read by the second plugin. The second plugin reads the table generated by the first plugin and summarizes the positions of different proteins in one plot. For each centriole, one protein (here the tubulin) is defined as the reference protein and its Y coordinates are shifted so its Y starting point is set to 0. The same shift is applied to the second protein to keep the correct distances between the 2 proteins. The Y coordinates values of both signals are then rescaled to the average length of the reference protein. For each protein, the average and side deviation of the Y starting position and of the Y ending position is calculated. A plot is then generated illustrating the average position of one protein to the other.

### Statistical analysis

The statistical significance of the difference between groups was determined using one-way ANOVA followed by Tukey’s post hoc test. All data are presented as an average +/- SEM as specified in the figure legends. Differences were considered significant when p<0.05. Results reported are from 2-3 independent biological replicates as noted in legends with reproducible findings each time.

## Acknowledgements

We would like to thank Carsten Janke for his generous gift of anti-poly-E antibodies. We acknowledge Mireille Bétermier and all members of the Bétermier lab for stimulating discussions. We thank B. Durand for critical reading of the manuscript. We thank Cindy Mathon for maintaining the Paramecium cell cycle and her great help in molecular biology and immunofluorescence staining and Pascaline Tirand for excellent technical assistance. The present work has benefited from Imagerie-Gif core facility supported by I’Agence Nationale de la Recherche (ANR-11-EQPX-0029/Morphoscope, ANR-10-INBS-04/FranceBioImaging; ANR-11-IDEX-0003-02/ Saclay Plant Sciences). This work has been founded by basal body anchoring in ciliogenesis: structure-function analysis: ANR-15-CE11-0002-01 to AMT. PLB was supported by PhD fellowships from Université Paris-Saclay (https://www.paris-saclay.fr/). This work has been supported by the Fondation ARC pour la recherche sur le cancer” to PLB ARCDOC4202003000178 and by the Swiss National Science Foundation (SNSF) PP00P3_187198 and by the European research Council ERC ACCENT StG 715289 attributed to Paul Guichard.

## Author contributions

AMT conceived, supervised, designed the project, and wrote the final manuscript with input from all authors. PLB, performed almost all experiments, image acquisition, processing and analyzed the data. MHL performed ultrastructure expansion on mammalian cells, images acquisition and analyzed and quantified all U-ExM data. LG performed the MNR and OFD1 siRNA experiments in RPE1. LG and KB performed the microtubule recruitment experiments induced by MNR; ML performed the electron microscopy of *Paramecium*. MLG wrote the U-ExM quantification ImageJ PlugIn. OR provided antibodies and cell lines. LL and GC performed mass spectrometry analysis. MT acquired with PLB the *Paramecium* U-ExM confocal images. FK, VH and PG performed formal analysis, review and editing.

## Conflict of interest

The authors have declared that no competing interests exist.

## Supplementary material

**Table S1: myc-BirA*-FOPNL BioID raw protein measurement**. MS/MS comparison of two biological replicates of the myc-BirA*-FOPNL with the myc-BirA*. High-confidence interactors were defined as those with q value <0.05 and max fold >5.

**Table S2: Primer, RNAi, siRNA and CEP90-GFP RNAi-resistant sequences.** All primers used to construct *Paramecium* CEP90-GFP and CEP90-GFP resistant plasmid as well as *CEP90a* and *CEP90b* gene silencing vectors were synthesized by Eurofins Genomics. The RNAi-resistant sequence of CEP90-GFP RNAi-resistant plasmid (red) was designed following the ciliated codon code and was synthesized by Integrated DNA technologies. siRNA sequences directed against CEP90 and FOPNL were synthesized by Dharmacon according to the manufacturer’s protocol. In both case a published single sequence were used (siRNA CEP90 #2 and siRNA FOPNL #1) as well as a combination of 4 sequences directed against the gene (smartpool).

**Figure S1:** FOPNL, CEP90 and OFD1 forms a complex in mammalian cells **A)** Heatmap of proteins identified in myc-BirA*-FOPNL + tet *versus* myc-BirA*+ tet, made with Qlucore. Proteins were filtered based on a p-value<0.05 and Fold Change>5, intensities were log_2_ transformed and were displayed as colours ranging from red to blue. Both rows and columns were clustered hierarchically. **B)** Protein-protein interaction network from differentially abundant proteins identified in FOPNL-BirA+tet *versus* BirA*+tet. Proteins were filtered based on a p-value<0.05 and Fold Change>5. The network was created using STRING DB representing a full STRING network at medium confidence (0.400) with disconnected nodes hidden. Lines of different thicknesses between nodes symbolize the Edge confidence from medium to the highest. The cluster encircled in purple indicates centriolar and centriolar satellite proteins. The one in green corresponds to nuclear proteins. **C)** Co-immunoprecipitation experiment made in HeLa Kyoto GFP-FOPNL. Cell extract was immunoprecipitated with mouse control or anti-GFP immunoglobulins. The input, the unbound fraction (FT) as well as the bound fraction (IP) were revealed with either rabbit polyclonal anti-GFP, CEP90 and OFD1 antibodies. Both CEP90 and OFD1 are co-immunoprecipitated by GFP antibodies, indicating a complex between FOPNL, OFD1 and CEP90.

**Figure S2: Localization of CEP90-GFP, OFD1-GFP and GFP-FOPNL by U-ExM. A**) EM on *Paramecium* cortex fragment showing a two-BB unit. During cortex purification, cilia shed above the TZ (white arrowhead). The unciliated BB (yellow arrowhead) remain unmodified. Scale bar= 100 nm. **B-E**) U-ExM performed on paramecia transformants expressing CEP90-GFP (**B**), OFD1-GFP (**C, E**) and GFP-FOPNL (**D**) show BB and cilia stained with tubulin antibody (magenta). Note that a space indicated by a white arrowhead shows the cilium breakage between the distal end of the TZ and the cilium. **B, C**) CEP90-GFP and OFD1-GFP signal are localized at the distal extremity of non-ciliated BB (yellow arrowhead) while GFP signal is slightly underneath the distal end of ciliated BB (white arrowhead). Scale bar= 250 nm. **F**) Schematic representation of OFD1 (blue square), FOPNL (yellow square) and CEP90 (green square) localization on ciliated and unciliated BB. Scale bar= 200 nm.

**Figure S3: CEP90 depletion phenotypes**. **A-C**): *Paramecium* were transformed with a construction expressing either WT CEP90-GFP or a RNAi-resistant CEP90-GFP. **A**) Quantification of BB GFP signal fluorescence on newly formed basal bodies (anterior ones) after 2 or 3 divisions upon CEP90 depletion. average ± SD is represented, ****p< 0.001 (one-way ANOVA followed by Tukey’s post hoc test), n ≥ 60 basal bodies from 2 independent experiments. **B**) Number of cell divisions in control, CEP90 and FOPNL depleted cells as well as in cells rescued with RNAi-resistant CEP90. Note that both CEP90 and FOPNL depleted cells died after 4 or 5 divisions. **C**) Quantification of the swimming speeds of non-injected paramecia (NI), paramecia expressing WT CEP90-GFP or RNAi-resistant CEP90-GFP after 48h under control condition or CEP90 RNAi condition. Each dot shows the average velocity of 1 cell, average± SD is represented, ****p< 0.001 (one-way ANOVA followed by Tukey’s post hoc test, n ≥ 120 cells per condition performed from 2 independent experiments).

**Figure S4: CEP90 depletion does not affect BB duplication**. **A)** Early stage of *Paramecium* division stained with poly-E antibodies (grey). On both sides of the future fission line, BB duplication takes place as observed by the presence of 3 to 4 BB in row instead of 1 or 2. Cells were observed after control (Ctrl^RNAi^) and CEP90 depletion (CEP90^RNAi^) during the first division. No significant duplication defects were observed after CEP90 depletion. Scale bars: 20µm and 2µm on the magnification insert **B**) After the first division, the posterior part of the cell show defective BB organization after CEP90 depletion, suggesting that the RNAi has been effective. The invariant field encircled in yellow on the figure, shows the usual 2 BB pattern at the cortical level, suggesting that BB duplication has occurred normally. Defective BB duplication will lead to only 1BB in this field. Sometimes additional BB are found but appear below the cortical surface (yellow arrowheads). Scale bars= 20µm.

**Figure S5: Recruitment of CEN2-GFP, CEP90-GFP, OFD1-GFP and GFP-FOPNL in Cen2, Cen3 and CEP90 depleted cells to newly formed BB**. **A**) Transformants expressing a GFP-tagged FOPNL were observed 2-3 divisions upon control (Ctrl) or CEP90 depletion. GFP-FOPNL fluorescence (green) was severely reduced on newly formed BB along with the expected BB pattern disorganization. Scale bar= 20µm. **B**) Dot plots showing the average percentage of GFP-FOPNL BB fluorescence after 2-3 divisions in either FOPNL or CEP90 RNAi (*n* ≥ 100 basal bodies per condition performed in 2 independent replicates). AU: arbitrary units. Error bars show SEM. Statistical significance was assessed by a one-way ANOVA followed by Tukey’s post hoc test p<0.0001 ****. **C, D**) *Paramecium* transformants expressing Cen2-GFP, CEP90 GFP or OFD1-GFP (green) have been stained for BB (poly-E antibodies, magenta) after 2-3 divisions upon Cen2 **(C)** or Cen3 **(D)** depletion. After Cen2 and Cen3 depletion, BB pattern disorganization is observed. **C)** The GFP fluorescence is not recruited to newly formed BB after Cen2 RNAi. **D)** By contrast, Cen3 depletion did not affect neither CEP90-GFP, OFD1-GFP and GFP-FOPNL recruitment to BB. Scale bars= 20µm and 2 µm (magnification).

**Figure S6: Localization of FOPNL, CEP90 and MNR**. **A**) HeLa-Kyoto cell lines expressing GFP-FOPNL (green) were stained for CEP90 (magenta). Confocal images show the colocalization of both proteins at centrosome and centriolar satellites. Scale bar= 5µm. **B**, **C**) Confocal images of duplicating U2OS cell, expanded and stained for tubulin (magenta) and CEP90 (green), OFD1 (cyan) or MNR (orange/red) showing the recruitment of the 3 proteins at early stages of procentriole assembly, slightly after the arrival of the tubulin (PC: procentriole, MC: mature centriole). Scale bars 200 nm. **D-F**) Confocal images of duplicating RPE1 cells, expanded and stained for tubulin (magenta) and CEP90 (green) showing the presence of CEP90 slightly underneath the distal end of both the mother (MC) and procentrioles (PC). Note that CEP90 is recruited at early stages of procentriole assembly just after the start of the tubulin (arrowheads) Scale bar= 200 nm. **G**) Confocal images of ciliated RPE1 cell, expanded and stained for tubulin (magenta) and CEP90 (green) showing the presence of CEP90 at the distal part of the BB templating the primary cilium (C). Please note that the daughter centriole (DC) is also positive for CEP90 as shown in the upright insert where the distal part of the DC is shown. Scale bar= 200 nm.

**Figure S7: Localization of CEP90, FOPNL, OFD1, CEP83, CEP89, CEP164 in FOPNL and CEP90 depleted cells. A**) Representative images of distal appendage proteins labelling after depletion of CEP90 (siRNA#2) or FOPNL (siRNA#2), stained with CEP90, FOPNL, OFD1, CEP83, CEP89, CEP164 antibodies (green). The centrosome is stained with GT335 (magenta). Control siRNA and CEP90 (siRNA#1) and FOPNL (siRNA#1) are found in Figure 6A. **B)** quantification of the fluorescence intensity of antibodies: CEP90, OFD1, MNR, FOPNL, CEP83 and CEP164 staining at the centrosome. AU: arbitrary units. All data are presented as average ± SEM. ****p< 0.001 (one-way ANOVA followed by Tukey’s post hoc test), n ≥ 100 centrosomes in 3 independent experiments.

**Figure S8: Effect of CEP90 and FOPNL silencing in primary cilia formation and centrobin localization. A, A’**) Serum-starved RPE1 during 24h in control (siCtrl) condition and upon CEP90 (**A**) or FOPNL (**A’**) depletion were stained for GT335 (magenta) and CEP90 (**A**, green) or FOPNL (**A’**, green) scale bars: 15µm. **B**) Percentage of ciliated cells were quantified in each condition. average ± SD is represented, ****p< 0.001 (one-way ANOVA followed by Tukey’s post hoc test, n ≥ 350 cells per condition performed in 3 independent replicates). **C**) The daughter centriolar protein Centrobin was localized in control (siCtrl), CEP90 and FOPNL depleted RPE1 cells. No significant differences are observed between control-depleted cells and CEP90 or FOPNL depleted cells. Scale bar= 5µm.

**Figure S9: Control experiments of the recruitment of endogenous CEP83 on microtubules by MNR-GFP and CEP90 overexpression. A)** U2OS cells overexpressing MNR-GFP stained for endogenous OFD1, CEP90 and CEP83 as well as the percentage of cells displaying endogenous proteins to either cytoplasm (Cyt) or microtubules (MT) Scale bars: 20µm. **B, D, F**) Overexpression of myc-FOPNL, CEP90 and OFD1 mcherry and their localization to the cytoplasm as well as the percentage of cells displaying the localization of proteins to either cytoplasm (Cyt) or microtubules (MT). **C, E, G**) Co-overexpression of myc-FOPNL, CEP90 and OFD1 mcherry with MNR-GFP. Note that the cytoplasmic localization of myc-FOPNL, CEP90 and OFD1 mcherry shift with MNR on the microtubules. Percentage of cells displaying the localization of proteins to either cytoplasm (Cyt) or microtubules (MT). Scale bars= 20µm

## Notes

### Competing Interest Statement

The authors have declared no competing interest.

