## Supplementary figures and images for "The evolutionary conserved complex CEP90, FOPNL and OFD1 specifies the future location of centriolar distal appendages, and promotes their assembly"

### Supplemental Figure 1

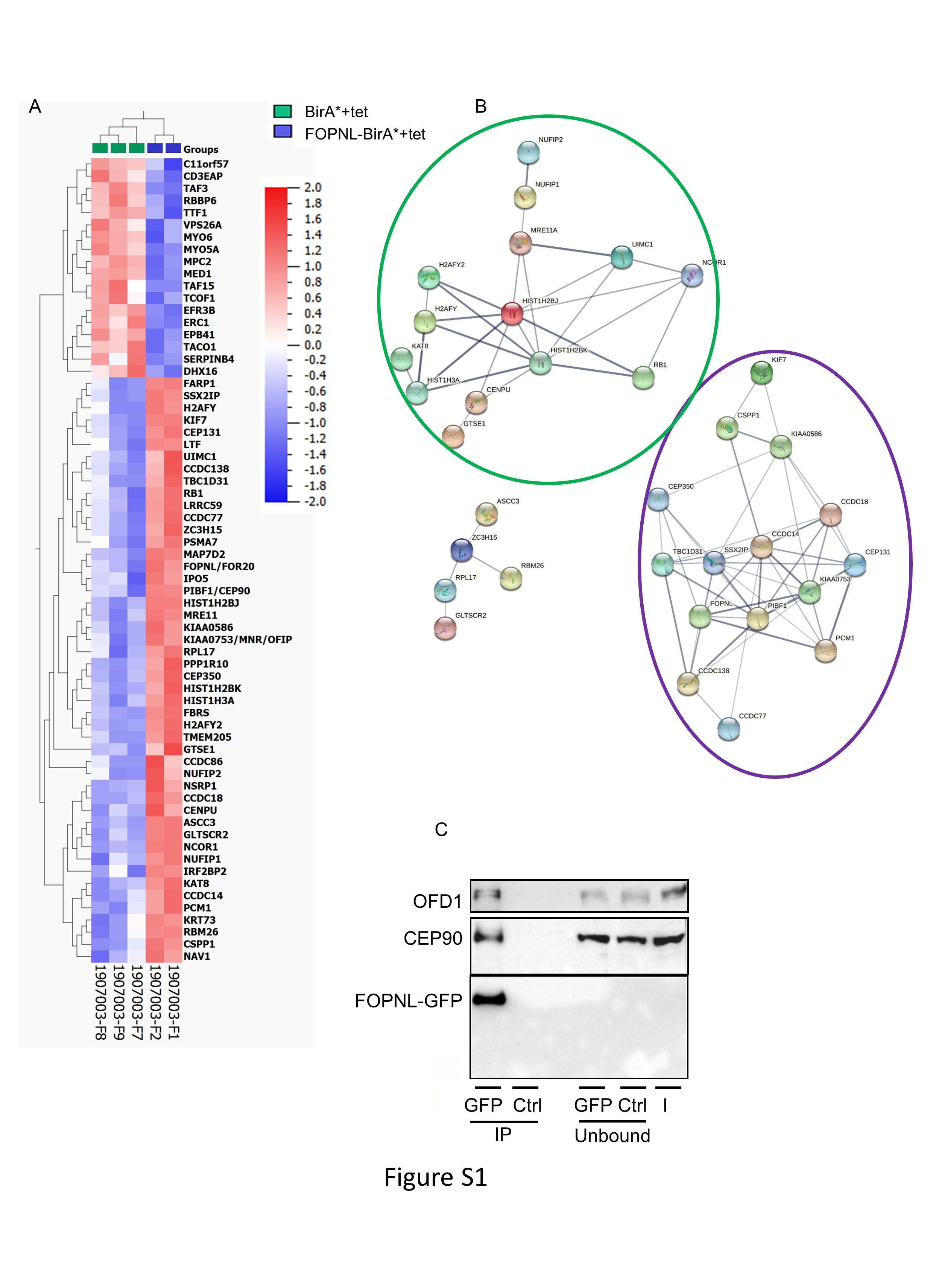

### Supplemental Figure 2

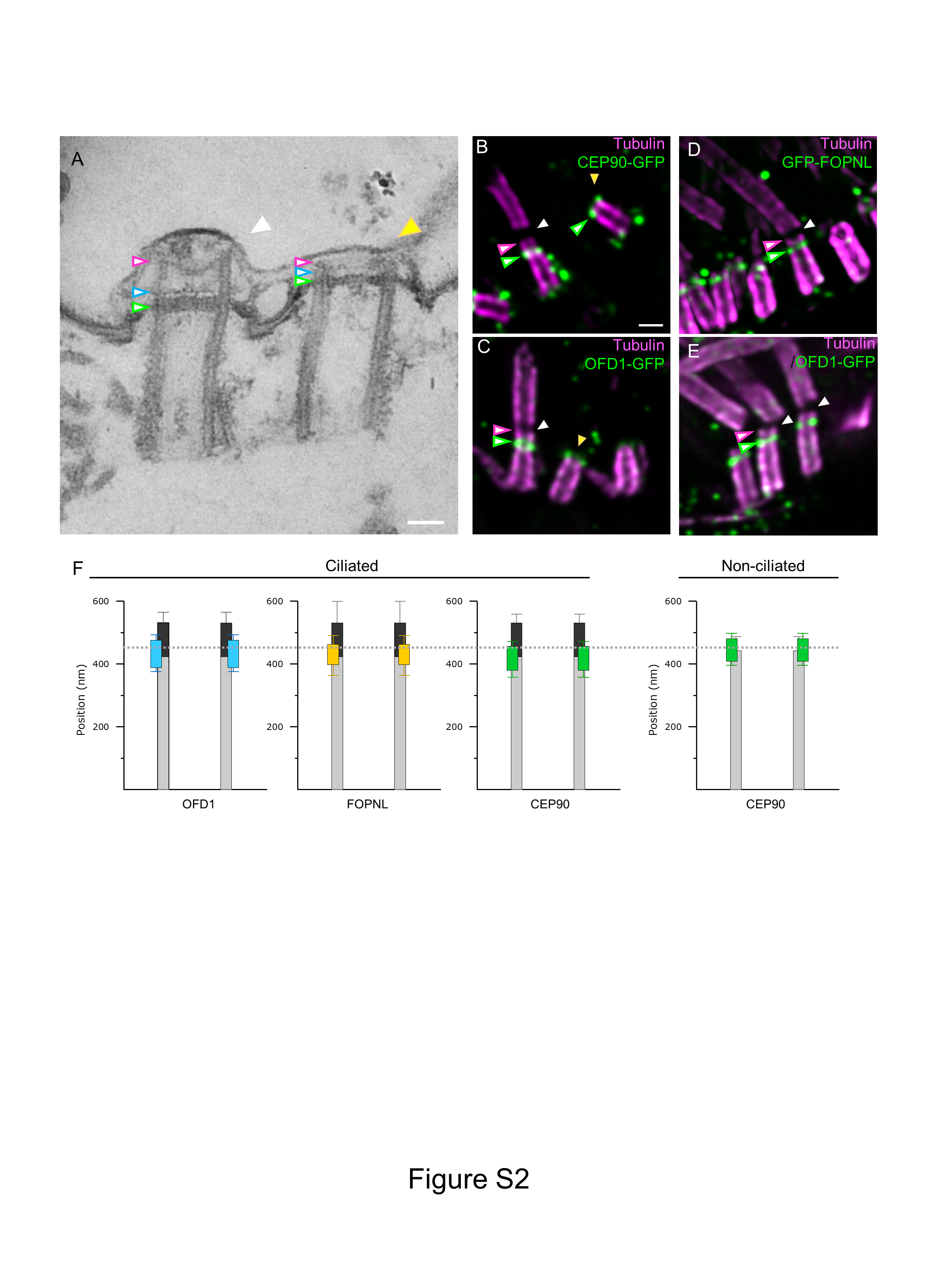

### Supplemental Figure 3

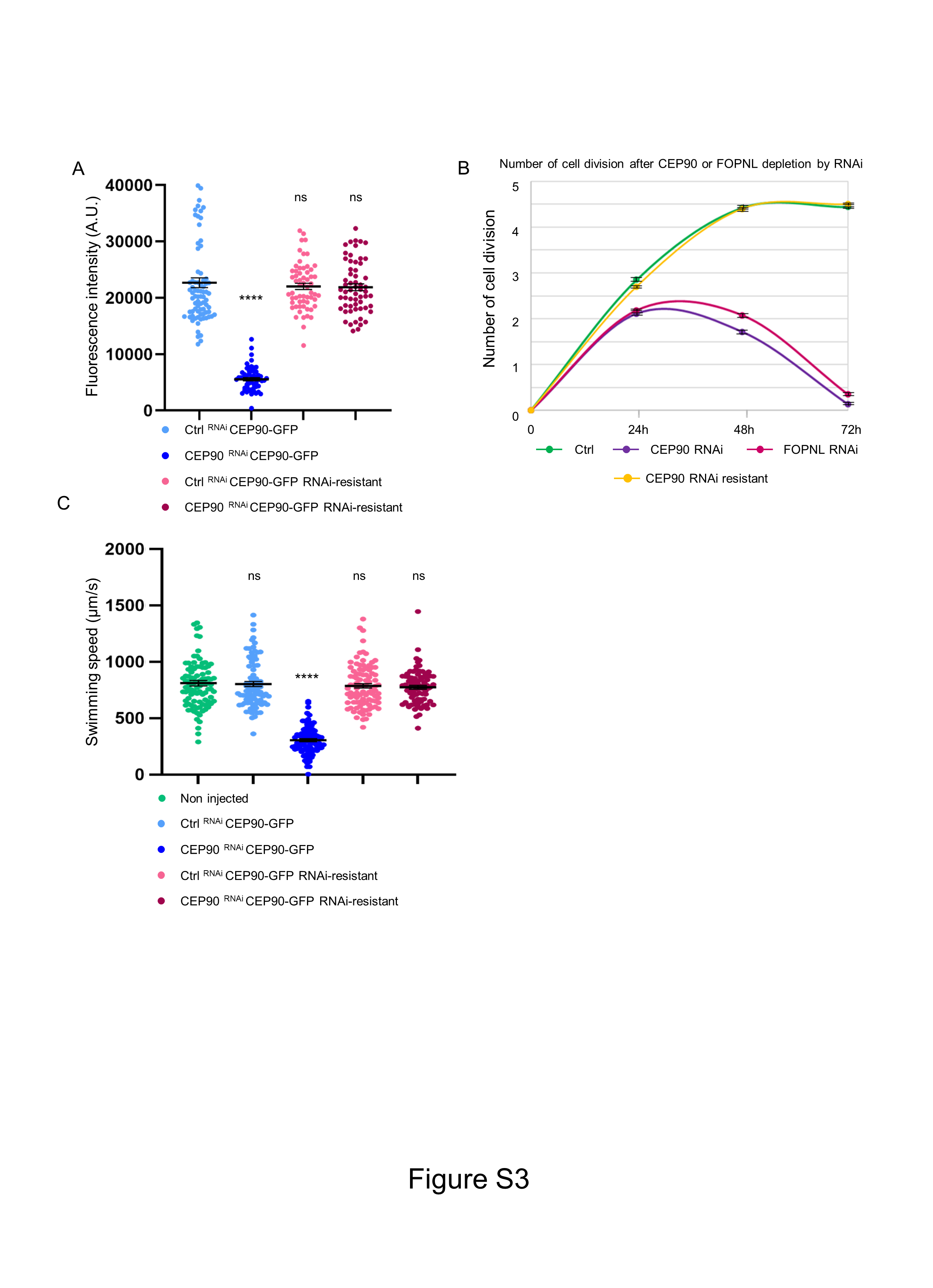

### Supplemental Figure 4

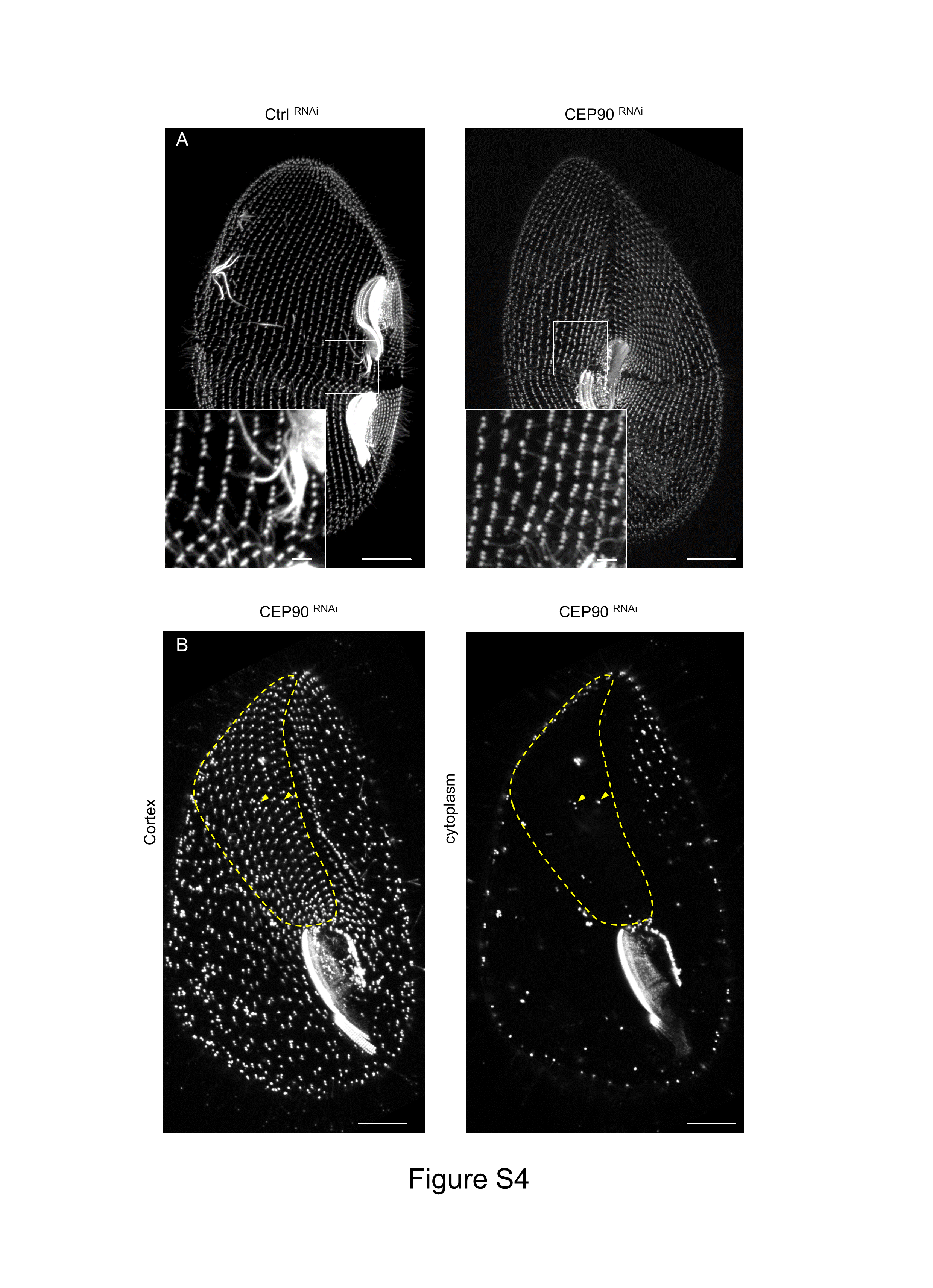

### Supplemental Figure 5

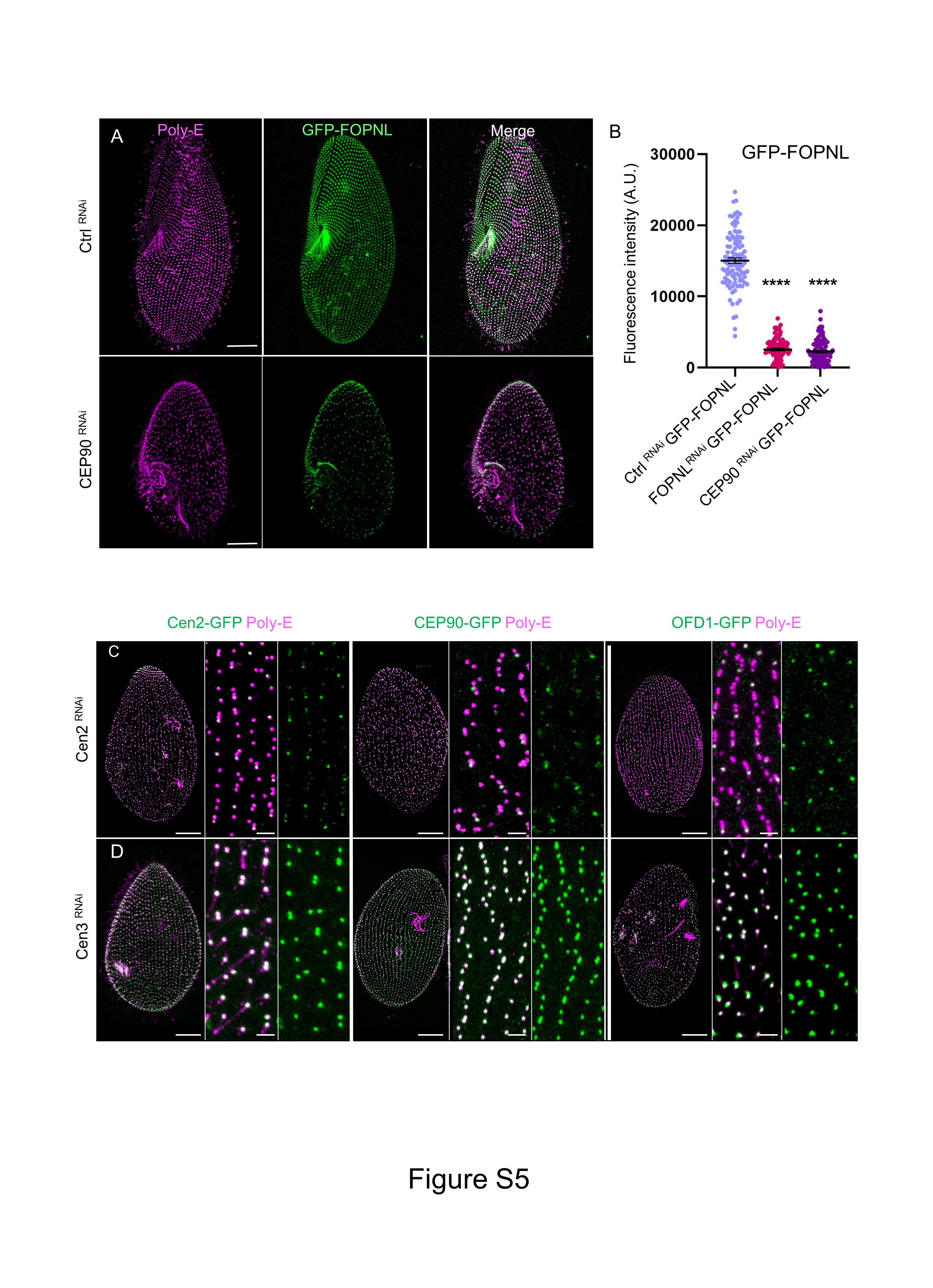

### Supplemental Figure 6

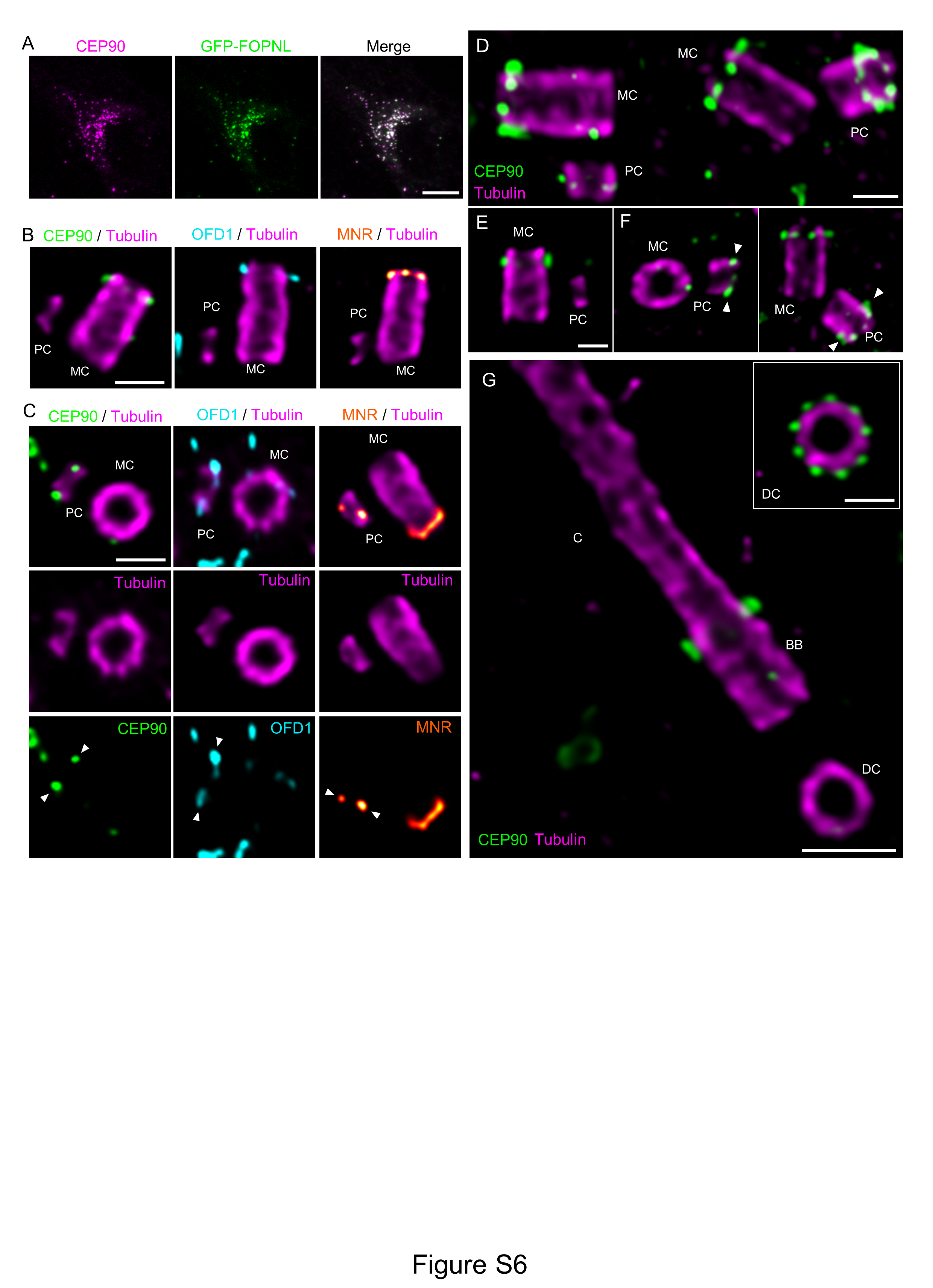

### Supplemental Figure 7

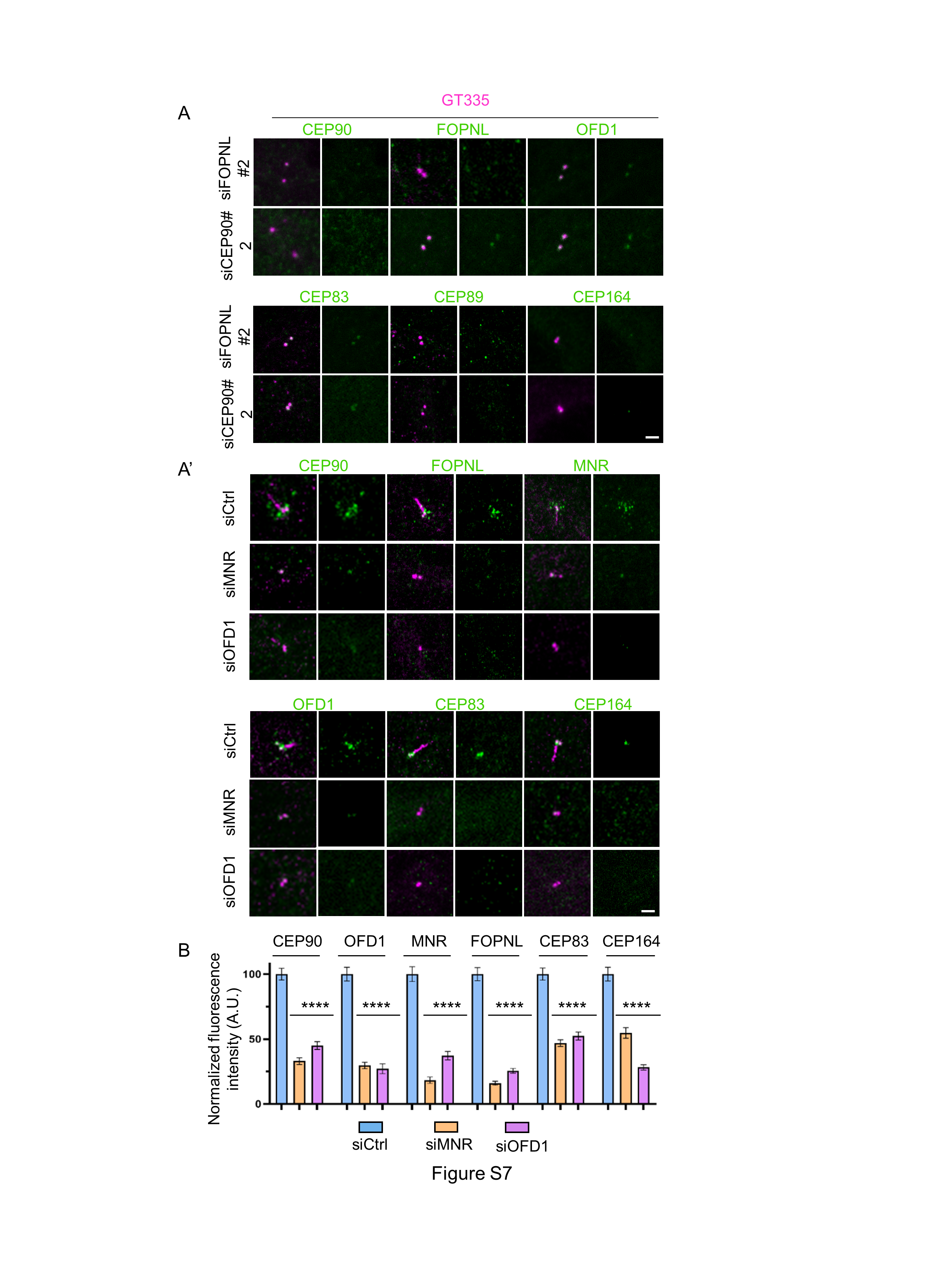

### Supplemental Figure 8

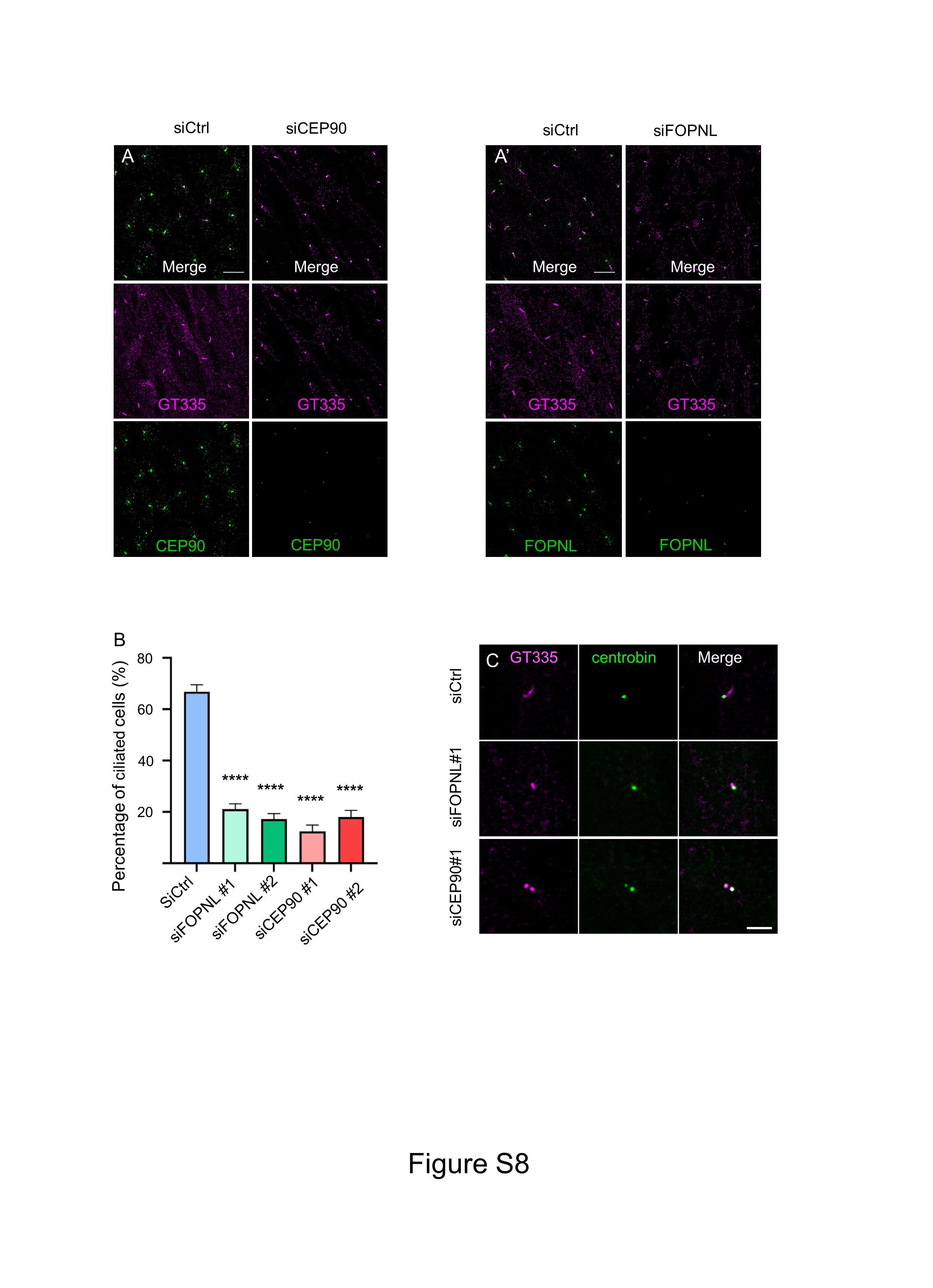

### Supplemental Figure 9

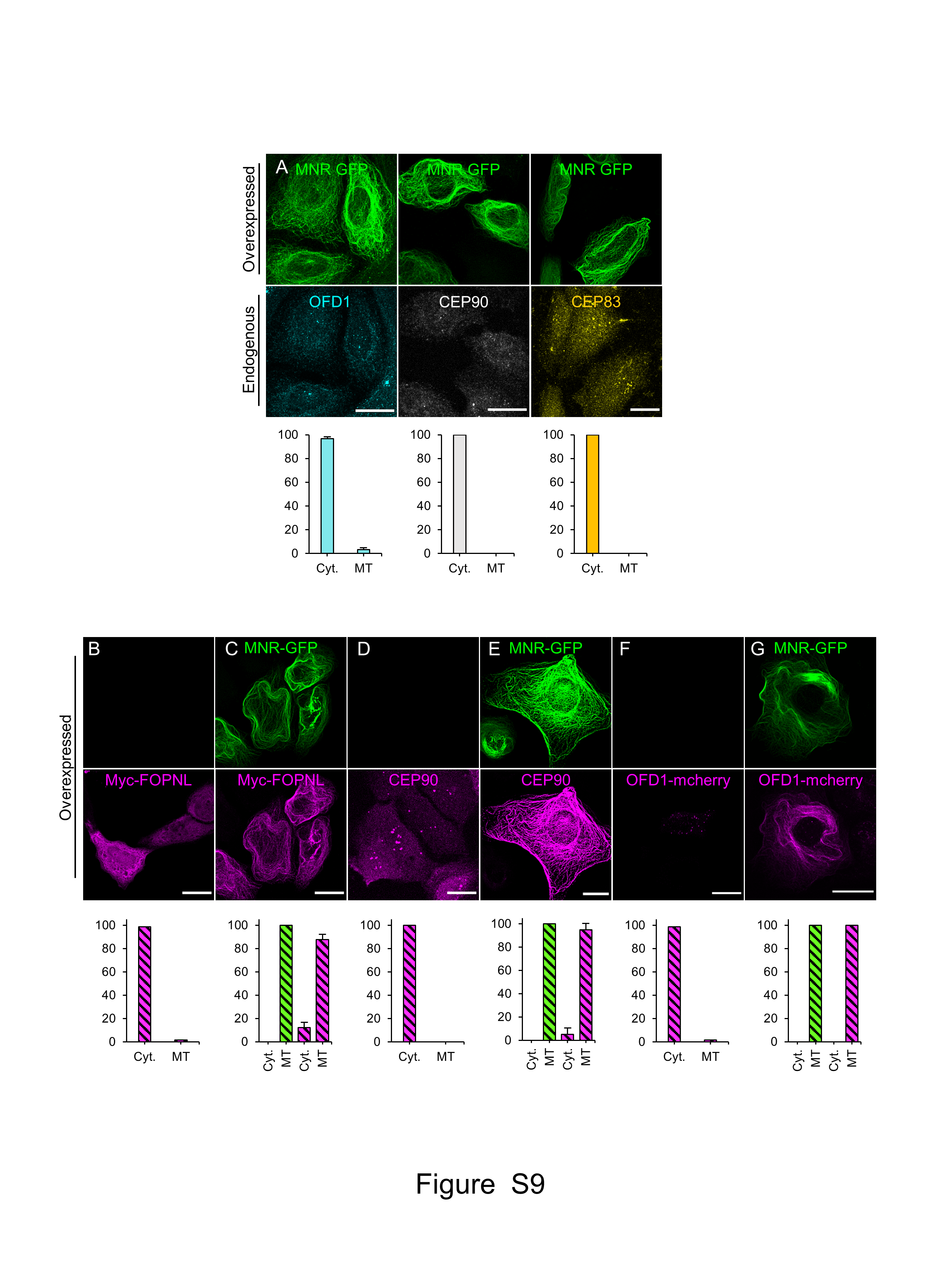
