## Supplemental Table 1 for "The evolutionary conserved complex CEP90, FOPNL and OFD1 specifies the future location of centriolar distal appendages, and promotes their assembly"

| Accession | Confidence score | Description | Gene | Mass | Peptide count | Unique peptides | p-value | Fold change | Groups Ids | for20-BirA+tet 1907003-F1 | BirA+tet 1907003-F9 | for20-BirA+tet 1907003-F2 | BirA+tet 1907003-F7 | BirA+tet 1907003-F8 |
|---|---|---|---|---|---|---|---|---|---|---|---|---|---|---|
| Q6ZUT1 | 67,05 | Uncharacterized protein C11orf57 | C11orf57 | 34,089 | 3 | 3 | 0,03360462 | 0,16797167 | | 2459,7 | 25312,4 | 8212,14 | 22180,4 | 34119,1 |
| O15446 | 1751,47 | DNA-directed RNA polymerase I subunit RPA34 | CD3EAP | 54,951 | 33 | 33 | 0,02480211 | 0,08785279 | | 49774,7 | 779839 | 103517 | 421026 | 1,66E+06 |
| Q5VWG9 | 1790,03 | Transcription initiation factor TFIID subunit 3 | TAF3 | 103,517 | 43 | 42 | 0,00467459 | 0,12848278 | | 90396,6 | 1,15E+06 | 109610 | 590966 | 686189 |
| Q7Z6E9 | 3188,3 | E3 ubiquitin-protein ligase RBBP6 | RBBP6 | 201,442 | 70 | 69 | 0,0173922 | 0,06080739 | | 38663,6 | 2,07E+06 | 102607 | 636105 | 844328 |
| Q15361 | 2230,86 | Transcription termination factor 1 | TTF1 | 102,987 | 42 | 42 | 0,0120319 | 0,16421238 | | 110606 | 1,21E+06 | 246821 | 1,01E+06 | 832216 |
| O75436 | 28,74 | Vacuolar protein sorting-associated protein 26A | VPS26A | 38,146 | 2 | 2 | 0,03607927 | 0,01825718 | | 283,994 | 7068,01 | 48,8272 | 1877,69 | 20217,8 |
| Q9UM54 | 44,82 | Unconventional myosin-VI | MYO6 | 149,596 | 3 | 3 | 0,01418116 | 0,19093625 | | 732,594 | 2687,78 | 346,233 | 2112,19 | 3232,63 |
| Q9Y4I1 | 93,39 | Unconventional myosin-Va | MYO5A | 215,269 | 4 | 4 | 0,00779616 | 0,03186501 | | 477,515 | 12114,3 | 312,587 | 6065,4 | 24257 |
| O95563 | 42,09 | Mitochondrial pyruvate carrier 2 | MPC2 | 14,27 | 2 | 2 | 0,00248825 | 0,00707438 | | 48,0314 | 6932,78 | 16,9828 | 3202,22 | 2964 |
| Q15648 | 1862,28 | Mediator of RNA polymerase II transcription subunit 1 | MED1 | 168,373 | 42 | 41 | 0,00133412 | 0,09684907 | | 59547,1 | 585973 | 40208,5 | 412806 | 533161 |
| Q92804 | 132,14 | TATA-binding protein-associated factor 2N | TAF15 | 61,793 | 3 | 3 | 0,03695844 | 0,00438169 | | 0 | 1287,14 | 0 | 24,4949 | 377,026 |
| Q13428 | 6757,45 | Treacle protein | TCOF1 | 152,015 | 155 | 155 | 0,03338429 | 0,10964112 | | 1,36E+06 | 1,65E+07 | 664099 | 4,07E+06 | 9,73E+06 |
| Q9Y2G0 | 41,27 | Protein EFR3 homolog B | EFR3B | 92,428 | 2 | 2 | 0,00209559 | 0,12504308 | | 404,6 | 2864,99 | 540,212 | 4509,4 | 4045,43 |
| Q8IUD2 | 625,25 | ELKS/Rab6-interacting/CAST family member 1 | ERC1 | 128,008 | 19 | 19 | 0,01245218 | 0,02357616 | | 1729,79 | 31255 | 2231,47 | 195037 | 94933,7 |
| P11171 | 96,68 | Protein 4.1 | EPB41 | 96,957 | 4 | 4 | 0,00614629 | 0,00084752 | | 4,65278 | 766,304 | 0 | 2964,21 | 7257,99 |
| Q9BSH4 | 36,42 | Translational activator of cytochrome c oxidase 1 | TACO1 | 32,457 | 2 | 2 | 0,0138146 | 0,01268087 | | 58,5538 | 1295,71 | 19,2958 | 8105,33 | 1773,38 |
| P48594 | 57,13 | Serpin B4 | SERPINB4 | 44,825 | 2 | 2 | 0,04323866 | 0,04444519 | | 18,938 | 88,8534 | 12,5337 | 937,957 | 499,79 |
| O60231 | 54,97 | Putative pre-mRNA-splicing factor ATP-dependent RNA helicase DHX16 | DHX16 | 119,189 | 3 | 3 | 0,02965721 | 0,09865307 | | 407,295 | 4844,93 | 733,001 | 12270,5 | 2857,86 |
| Q9Y4F1 | 48,8 | FERM, RhoGEF and pleckstrin domain-containing protein 1 | FARP1 | 118,559 | 2 | 2 | 0,01505139 | 6,04822813 | | 6843,86 | 770,747 | 6375,24 | 897,746 | 1882,56 |
| Q9Y2D8 | 202,89 | Afadin- and alpha-actinin-binding protein | SSX2IP | 71,191 | 7 | 6 | 0,00949265 | 18,2411837 | | 30540,4 | 1198,41 | 39850,1 | 1469,64 | 3971,73 |
| O75367 | 442,18 | Core histone macro-H2A.1 | H2AFY | 39,592 | 8 | 7 | 0,02704455 | 7,22323884 | | 1,21E+06 | 130705 | 1,55E+06 | 130736 | 397916 |
| Q2M1P5 | 102,28 | Kinesin-like protein KIF7 | KIF7 | 150,495 | 4 | 3 | 0,00807763 | 50,153359 | | 5834,75 | 85,4212 | 5672,45 | 60,1175 | 293,916 |
| Q9UPN4 | 734,35 | Centrosomal protein of 131 kDa | CEP131 | 122,075 | 15 | 15 | 0,01359155 | 17,6582979 | | 165735 | 6364,08 | 110409 | 4218,37 | 16745,6 |
| P02788 | 111,19 | Lactotransferrin | LTF | 78,132 | 6 | 6 | 0,03378333 | 6,09722837 | | 19056,6 | 2681,32 | 19748,1 | 1837,54 | 6536,86 |
| Q96RL1 | 180,66 | BRCA1-A complex subunit RAP80 | UIMC1 | 79,678 | 8 | 8 | 0,03781041 | 7,41178388 | | 50021,3 | 3962,14 | 16545 | 2299,44 | 6418,19 |
| Q96M89 | 338,97 | Coiled-coil domain-containing protein 138 | CCDC138 | 76,171 | 9 | 9 | 0,02516265 | 20,4977784 | | 157083 | 3298,62 | 38360,3 | 2098,48 | 7846,2 |
| Q96DN5 | 117,44 | TBC1 domain family member 31 | TBC1D31 | 124,111 | 4 | 4 | 0,03012343 | 7,87519379 | | 41072,5 | 2513,38 | 17974,9 | 2426,18 | 6735,39 |
| P06400 | 72,57 | Retinoblastoma-associated protein | RB1 | 106,092 | 2 | 2 | 0,03147326 | 56,1256136 | | 33551,6 | 391,016 | 11903,2 | 91,5071 | 1261,63 |
| Q96AG4 | 448,15 | Leucine-rich repeat-containing protein 59 | LRRC59 | 34,909 | 10 | 10 | 0,02113737 | 6,1892979 | | 797862 | 115700 | 563576 | 62425,1 | 176077 |
| Q9BR77 | 50,28 | Coiled-coil domain-containing protein 77 | CCDC77 | 57,45 | 4 | 4 | 0,01975279 | 21,1960996 | | 13962,1 | 408,959 | 6117,12 | 203,131 | 997,756 |
| Q8WU90 | 143,87 | Zinc finger CCCH domain-containing protein 15 | ZC3H15 | 48,573 | 5 | 5 | 0,02287182 | 5,86262707 | | 57952,2 | 6818,87 | 30366,1 | 4709,6 | 11408,1 |
| O14818 | 102,9 | Proteasome subunit alpha type-7 | PSMA7 | 27,87 | 3 | 3 | 0,04049976 | 5,69755927 | | 23694,8 | 2940,02 | 15950,1 | 1955,71 | 6908,85 |
| Q96T17 | 139,92 | MAP7 domain-containing protein 2 | MAP7D2 | 81,915 | 5 | 5 | 0,00347276 | 12,6879338 | | 15784,4 | 1559,56 | 18112,1 | 848,197 | 1789,06 |
| Q96NB1 | 655,78 | LisH domain-containing protein FOPNL | FOPNL/FOR2( | 19,765 | 11 | 11 | 0,01453903 | 73,240904 | | 650665 | 16468,2 | 1,54E+06 | 4438,31 | 34864,5 |
| O00410 | 22,19 | Importin-5 | IPO5 | 123,55 | 2 | 2 | 0,03687303 | 26,1411726 | | 2837,84 | 318,764 | 5102,61 | 37,3864 | 258,829 |
| Q8WXW3 | 593,59 | Progesterone-induced-blocking factor 1 | PIBF1/CEP90 | 89,75 | 17 | 17 | 0,01978298 | 26,9834882 | | 198501 | 11801,7 | 212056 | 2503,17 | 14879,6 |
| P06899 | 73,2 | Histone H2B type 1-J | HIST1H2BJ | 13,896 | 2 | 2 | 0,00179869 | 6,00162879 | | 1,14E+06 | 144082 | 1,10E+06 | 206696 | 218742 |
| P49959 | 849,76 | Double-strand break repair protein MRE11 | MRE11 | 80,543 | 24 | 24 | 0,00643677 | 5,24805818 | | 1,16E+06 | 164325 | 1,32E+06 | 300239 | 266489 |
| Q9BVV6 | 44,38 | Protein TALPID3 | KIAA0586 | 169,201 | 2 | 2 | 0,00871489 | 53,6774397 | | 2641,79 | 34,4187 | 6044,72 | 75,176 | 159,464 |
| Q2KHM9 | 1005,83 | Protein moonraker | KIAA0753/MN | 109,339 | 30 | 30 | 0,02024527 | 111,817687 | | 492067 | 1590,2 | 993389 | 6431,72 | 23900,4 |
| P18621 | 244,24 | 60S ribosomal protein L17 | RPL17 | 21,383 | 6 | 6 | 0,02830328 | 10,3889412 | | 666100 | 23638 | 427787 | 53561,1 | 107146 |
| Q96QC0 | 451,24 | Serine/threonine-protein phosphatase 1 regulatory subunit 10 | PPP1R10 | 98,996 | 15 | 15 | 0,00687355 | 9,47347435 | | 389828 | 21620,5 | 179335 | 34031 | 29548,7 |
| Q5VT06 | 1377,34 | Centrosome-associated protein 350 | CEP350 | 350,716 | 42 | 42 | 0,00498279 | 16,4401295 | | 256238 | 7369,33 | 120764 | 14067,4 | 11817,2 |
| O60814 | 618,13 | Histone H2B type 1-K | HIST1H2BK | 13,882 | 8 | 8 | 0,00867993 | 6,76859118 | | 2,68E+07 | 2,19E+06 | 1,35E+07 | 2,87E+06 | 3,53E+06 |
| P68431 | 62,83 | Histone H3.1 | HIST1H3A | 15,394 | 3 | 3 | 0,00872919 | 28,2471528 | | 241777 | 2855,96 | 114195 | 8871,08 | 8034,18 |
| Q9HAH7 | 121,41 | Probable fibrosin-1 | FBRS | 48,358 | 3 | 3 | 0,00255158 | 15,0097307 | | 94754,7 | 4629,49 | 63342,8 | 3972,01 | 7477,99 |
| Q9P0M6 | 291,6 | Core histone macro-H2A.2 | H2AFY2 | 40,033 | 9 | 9 | 0,00174913 | 12,9653627 | | 477950 | 24420,9 | 304288 | 27434,7 | 37982,4 |
| Q6UW68 | 60,46 | Transmembrane protein 205 | TMEM205 | 21,184 | 2 | 2 | 0,00720887 | 8,60639127 | | 57670,6 | 4184,21 | 33482,5 | 4176,49 | 7616,76 |
| Q9NYZ3 | 41,39 | G2 and S phase-expressed protein 1 | GTSE1 | 76,598 | 2 | 2 | 0,03185181 | 18,7547503 | | 7848,5 | 233,825 | 1238,8 | 98,0396 | 200,473 |
| Q9H6F5 | 397,94 | Coiled-coil domain-containing protein 86 | CCDC86 | 40,211 | 12 | 12 | 0,03939868 | 7,42632012 | | 39836,9 | 7555,1 | 134337 | 7469,81 | 16937,5 |
| Q7Z417 | 529,88 | Nuclear fragile X mental retardation-interacting protein 2 | NUFIP2 | 76,075 | 13 | 13 | 0,04342793 | 6,43184819 | | 63930,4 | 11353,8 | 169511 | 12124,8 | 30798,6 |
| Q9H0G5 | 100,4 | Nuclear speckle splicing regulatory protein 1 | NSRP1 | 66,35 | 4 | 4 | 0,00692071 | 16,6891404 | | 3296,8 | 350,099 | 9755,72 | 394,891 | 283,83 |
| Q5T9S5 | 313 | Coiled-coil domain-containing protein 18 | CCDC18 | 168,857 | 10 | 10 | 0,0019736 | 54,9892378 | | 54109,4 | 1263,31 | 128868 | 2130,73 | 1300,93 |
| Q71F23 | 214,13 | Centromere protein U | CENPU | 47,493 | 7 | 7 | 0,01644822 | 6,87021484 | | 13348,1 | 4183,32 | 30085,7 | 2786,17 | 2129,27 |
| Q8N3C0 | 107,52 | Activating signal cointegrator 1 complex subunit 3 | ASCC3 | 251,301 | 3 | 3 | 0,00110979 | 19,7408933 | | 93607,4 | 6471,67 | 84018,9 | 3732,17 | 3753,65 |
| Q9NZM5 | 163,68 | Glioma tumor suppressor candidate region gene 2 protein | GLTSCR2 | 54,356 | 4 | 4 | 0,00447235 | 5,23704304 | | 53106,6 | 13459,3 | 49462,6 | 8943,06 | 7787,04 |
| O75376 | 98,94 | Nuclear receptor corepressor 1 | NCOR1 | 270,044 | 4 | 4 | 0,00047959 | 7,81809772 | | 22503,9 | 3226,17 | 21918,9 | 3018,6 | 2354,06 |
| Q9UHK0 | 103,85 | Nuclear fragile X mental retardation-interacting protein 1 | NUFIP1 | 56,264 | 4 | 4 | 0,01919252 | 5,4726506 | | 5722,88 | 1489,28 | 4714,45 | 1010,74 | 568,011 |
| Q7Z5L9 | 34,07 | Interferon regulatory factor 2-binding protein 2 | IRF2BP2 | 60,987 | 2 | 2 | 0,03091246 | 27,9868248 | | 4940,95 | 689,411 | 5965,43 | 71,5304 | 148,03 |
| Q9H7Z6 | 73,94 | Histone acetyltransferase KAT8 | KAT8 | 52,37 | 2 | 2 | 0,00437548 | 11,0511586 | | 121855 | 9814,32 | 81431 | 12292,4 | 6070,54 |
| Q49A88 | 203,62 | Coiled-coil domain-containing protein 14 | CCDC14 | 106,236 | 5 | 5 | 0,01411943 | 31,5983576 | | 42565,8 | 700,497 | 17715,1 | 2072,02 | 452,183 |
| Q15154 | 3282,87 | Pericentriolar material 1 protein | PCM1 | 228,392 | 75 | 75 | 0,01922071 | 18,2449583 | | 1,90E+06 | 38735,6 | 931762 | 171076 | 58602,7 |
| Q86Y46 | 144,94 | Keratin, type II cytoskeletal 73 | KRT73 | 58,887 | 2 | 2 | 0,03710963 | 5,34510267 | | 158148 | 25930,4 | 176355 | 62704,9 | 18758,6 |
| Q5T8P6 | 169,57 | RNA-binding protein 26 | RBM26 | 113,527 | 4 | 4 | 0,02715926 | 5,97540095 | | 23247,5 | 3130,68 | 22217,9 | 7395,59 | 2376,34 |
| Q1MSJ5 | 107,92 | Centrosome and spindle pole-associated protein 1 | CSPP1 | 145,433 | 6 | 6 | 0,01251096 | 262,363165 | | 20706,9 | 63,0117 | 46091,2 | 540,803 | 47,9105 |
| Q8NEY1 | 89,07 | Neuron navigator 1 | NAV1 | 202,347 | 3 | 3 | 0,03116214 | 6,01483357 | | 2373,69 | 489,605 | 3563,37 | 869,497 | 265,548 |
