## Supplemental Table 2 for "The evolutionary conserved complex CEP90, FOPNL and OFD1 specifies the future location of centriolar distal appendages, and promotes their assembly"

| Table S2 | Primers, RNAI, siRNA and CEP90-GFP RNAI resistant sequences | |
|---|---|---|
| | **Primer Paramecium** | |
| **Constructs** | **Primers** | |
| **CEP90a-GFP** | CEP90a sens | caaatcattaattaaataaatcaactagtATGAGTACGAAGAGAAAAGACATCATAAAAC |
| | CEP90a anti-sens | CTTCTCCTTTAGACATgcgtacctcgagTCCTCCTCCTTTATTCTGTTTTTGTTTC |
| **CEP90a-GFP RNAi resistan** | CEP90a-GFP RNAi resistant sens A | GGATAGAGTATAGAATATCTATGAAGATACATTGGCTAATTTGAAAGCCACAAAACATGAAAACG |
| | CEP90a-GFP RNAi resistant antisens A | CGTTTTCATGTTTTGTGGCTTTCAAATTAGCCAATGTATCTTCATAGATATTCTATACTCTATCC |
| | CEP90a-GFP RNAi resistant sens B | GGATAAAACCCACTAACCTCACTCCTACCTCGTCGCCCATATTGAGGAAAAGGATAGAGAAATTTTG |
| | CEP90a-GFP RNAi resistant antisens B | CAAAATTTCTCTATCCTTTTCCTCAATATGGGCGACGAGGTAGGAGTGAGGTTAGTGGGTTTTATCC |
| **RNAi CEP90** | CEP90a sens | CCATTTAAAAAGGAAGGGGAATTTTAGAAACTTTGGCCAATCTTAAAGCTACAAAGCATG |
| | CEP90a antisens | CACCGCGGTGGCGGCCGCTCTAGAACTAGTTACGAGATAGGAATGActaaaataaaag |
| | CEP90b sens | CCATTTAAAAAGGAAGGGGAATTTTAGAAACTTTGGCCAATCTTAAAGCTACAAAGCATG |
| | CEP90b antisens | CATGCTTTGTAGCTTTAAGATTGGCCAAAGTTTCTAAAATTCCCCTTCCTTTTTAAATGG |
| | **RNAi *Paramecium*** | |
| **RNAi CEP90a** | | CTTTGGCCAATCTTAAAGCTACAAAGCATGAAAATGAAATGTTAAGGGAGAAGATCAACGTGTTGAAGGCAGAGTATTATAAGTGTTAGGTAGAGGCTAAGGAATAGATGAGCAGCATAAATGCTTAATTGCAAGTTGCAAAAGAACAATTAACTAATTATGAAGGAATTGAAAAAGAAATTGATGATGC<br>CATAATGAAATCAGCAGGCAATGAATATGATGCAATGAATCCTTTATTGGGATTCATGGGCACTGTACCTACATCTTCAAAAAGAAGGATTCAACAGGCATTAAATCTCGCTTAGAGATTGCAAGCAAAATAAAGAGAATCAGAGGAATTGCAAAGACAATTAAGAGATAAGTCATAAGAGATTGAGAGA<br>CTATAGGAAGAGACCAAATTATAAAGAGATGTACTTGACAAAACACATTAACCgtaaatattcattaaatatcttttattttagTCATTCCTATCTCGTA |
| **RNAi CEP90b** | | GATTCTGTATCCATAGCTGATATTGAGAATTTTTAATAGAAGAGACATAATAGAGGAGGGGATACTCACGACACAGTCAATATGAAAAATgtattataattgattcaataatagAATTATTCTTCTGTAGTGGGCATGCAAGAGAAATAAATAGCATTTCTAAATGCAGAAATAAAGCGGATTCAAGCAGCTAACGATGCT<br>GATAGACAAAGGATTTTTGCTAATCATAGAATTGAAACTGATAAGCTTTTAAGTGAAATCAAACAACTTAAGTCTAAGATGATATATTAAGAATCTCAATTGCCAGGAATAAGGGAGGCTTTGGCATAAGTCAGAGACATGTTAGCTGGGTTAGTTCCAGAGGTTGTTTATTTGAGATTGAGAGATATGAA<br>TGAGAAGGAGATGCCCATCTAAGATTGGGTGCTGGTTTAGGTTTGGGAGATTGTGTATCCATTTAAAAAGGAAGGGGAATTTTAGAAA |
| | **RNAi resistant CEP90-GFP** | |
| **CEP90a-GFP RNAi resistant** | | ATGAGTACGAAGAGAAAAGACATCATAAAACTTATGAAATCGATGAATATAGATCCAACCACATTTTCTGAGGAAGATATAAGATAATTTGAATAAGGCTAAGTGATGATGAGTGACAGTGAATCAGATAGTATGAGGGATTCGGTATCCATAGCTGATATTGAGAATTATCAATAGACTAAACATAACA<br>GAGGGGGGAACACTCATGATACAGTCAATATGAAAAATGTATTATTGTAATCGATCCATTTTTAGAATTATTCATCCGTTGTGGGCATGCAGGAAAAATAAATAGCTTTTCTAAATGCTGAGATCAAAAGGATCCAAGCAGCCAATGACGCTGATAGACAAAGAGTTTTTGCTAATCATCGAATTGAAACA<br>GATAAACTTTTAAGTGAAATCAAATAACTAAAGTCTAAGATGATCTATCAAGAGTCTCAATTACCAGGCATAAGAGAAGCTTTGGCTTAAGTTAGAGATATGCTTGCTGGATTGGTCCCAGAGGCTGTTTATCTGAGGTTGCGAGATATGAATGAGAAGGAAATGCCCATTCAAGATTGGGTATTGGTTTA<br>GGTTTGGGAAATTGTGTATCCATTCAAAAAGGAGGGGGAGTTTTAGAAAAAGGAAATATTGGCTTTAAGGGAAGAACTCAGATAGATAACAGAGAAGCATTAATTCTTATTGAATGATTTGGAACATGGCTAAAAGATGATGGTAGACAGAGAAGAGGATATTAGAAGACATAAATTAAATTATGATAAT<br>GCTAGAAAGGCTTTGGAGTTAGAATTAACAAAATCTTAAGAGGAATTAGCCCTGTTGAGAGAGAAGGGAGCAAGATTTGATGAATTAAGTAGGGATTATAAGAGATTGGAGTAAGAGAAGTTTTTATTGGAAGAGAAGGCTGGTTTCTATCAAAATAATTAAGCCGGAGACAAGGCAGGTGATTAATAC<br>AAGGCATAGGATGACGCCAAGAGGAAGGGAGACCTTTTGCAATAAGACAAGGAATATTTGACTAAGGAGAATATTTAACTTTTGGAGAAGGTTAAGAGATTGGAGGACAAATTAGATAGGACAGAGAAGGAGTATTTGGAAGCAAAAAACTAAGCTTAAGAATATCTATTTTAATTACTCAATTCTAAAA<br>CAGATTAGACTCAGGCTTATGAAAAAAGAATTTACTCAGAAATTGCAGATCTTAAAGAGAAGCATCATCATGAATTGGAAATTGCCAAATAGAATTTAGTTGATATTTATGAGACTCAAATCAAATTCTTGAAGGATGCAAAGGAGGAAATTTAACTTCGAAAAGATAATCTAGAAGGGCAAGTGAAGGA<br>AAAATCAACTTTATATGATCATTTGCTAATTGATTATAGAAACTTATAAAGAAAATTAGATGGGGACTTATCTGAATAGAGAATTCAATTAAGATTGAAGGTGGAAGAGTTGGATAGAGTATAGAATATCTATGAAGATA**CATTGGCTAATTTGAAAGCCACAAAACATGAAAACGAGATGTTGAGGG<br>AAAAAATCAATGTCTTAAAAGCCGAATATTATAAATGCTAAGTCGAGGCCAAGGAATAAATGTCAAGCATCAACGCTTAACTTTAAGTCGCCAAAGAATAATTGACTAACTATGAAGGAATCGAAAAGGAAATCGATGATGCTATTATGAAGTCAGCCGGAAACGAATACGATGCTATGA<br>ACCCTTTATTGGGATTCATGGGAACAGTCCCAACATCATCTAAAAGGAGAATTCAATAAGCATTGAACCTCGCCTAAAGACTTTAAGCAAAGCAAAGAGAAAGCGAAGAGCTTTAAAGACAATTGAGAGACAAATCACAAGAATCGAGAGACTTTAAGAGGAAACCAAATTACAAAGA<br>GATGTCTTGGATAAAACCCACTAACCTCACTCCTACCTCGTC**ATGAGTACGAAGAGAAAAGACATCATAAAACTTATGAAATCGATGAATATAGATCCAACCACATTTTCTGAGGAAGATATAAGATAATTTGAATAAGGCTAAGTGATGATGAGTGACAGTGAATCAGATAGTATGAGGGATTCGGT<br>ATCCATAGCTGATATTGAGAATTATCAATAGACTAAACATAACAGAGGGGGGAACACTCATGATACAGTCAATATGAAAAATGTATTATTGTAATCGATCCATTTTTAGAATTATTCATCCGTTGTGGGCATGCAGGAAAAATAAATAGCTTTTCTAAATGCTGAGATCAAAAGGATCCAAGCAGCCAATG<br>ACGCTGATAGACAAAGAGTTTTTGCTAATCATCGAATTGAAACAGATAAACTTTTAAGTGAAATCAAATAACTAAAGTCTAAGATGATCTATCAAGAGTCTCAATTACCAGGCATAAGAGAAGCTTTGGCTTAAGTTAGAGATATGCTTGCTGGATTGGTCCCAGAGGCTGTTTATCTGAGGTTGCGAGAT<br>ATGAATGAGAAGGAAATGCCCATTCAAGATTGGGTATTGGTTTAGGTTTGGGAAATTGTGTATCCATTCAAAAAGGAGGGGGAGTTTTAGAAAAAGGAAATATTGGCTTTAAGGGAAGAACTCAGATAGATAACAGAGAAGCATTAATTCTTATTGAATGATTTGGAACATGGCTAAAAGATGATGGTAG<br>ACAGAGAAGAGGATATTAGAAGACATAAATTAAATTATGATAATGCTAGAAAGGCTTTGGAGTTAGAATTAACAAAATCTTAAGAGGAATTAGCCCTGTTGAGAGAGAAGGGAGCAAGATTTGATGAATTAAGTAGGGATTATAAGAGATTGGAGTAAGAGAAGTTTTTATTGGAAGAGAAGGCTGGTT<br>TCTATCAAAATAATTAAGCCGGAGACAAGGCAGGTGATTAATACAAGGCATAGGATGACGCCAAGAGGAAGGGAGACCTTTTGCAATAAGACAAGGAATATTTGACTAAGGAGAATATTTAACTTTTGGAGAAGGTTAAGAGATTGGAGGACAAATTAGATAGGACAGAGAAGGAGTATTTGGAAGCAA<br>AAAACTAAGCTTAAGAATATCTATTTTAATTACTCAATTCTAAAACAGATTAGACTCAGGCTTATGAAAAAAGAATTTACTCAGAAATTGCAGATCTTAAAGAGAAGCATCATCATGAATTGGAAATTGCCAAATAGAATTTAGTTGATATTTATGAGACTCAAATCAAATTCTTGAAGGATGCAAAGGAG<br>GAAATTTAACTTCGAAAAGATAATCTAGAAGGGCAAGTGAAGGAAAAATCAACTTTATATGATCATTTGCTAATTGATTATAGAAACTTATAAAGAAAATTAGATGGGGACTTATCTGAATAGAGAATTCAATTAAGATTGAAGGTGGAAGAGTTGGATAGAGTATAGAATATCTATGAAGATACATTGG<br>CTAATTTGAAAGCCACAAAACATGAAAACGAGATGTTGAGGGAAAAAATCAATGTCTTAAAAGCCGAATATTATAAATGCTAAGTCGAGGCCAAGGAATAAATGTCAAGCATCAACGCTTAACTTTAAGTCGCCAAAGAATAATTGACTAACTATGAAGGAATCGAAAAGGAAATCGATGATGCTATTAT<br>GAAGTCAGCCGGAAACGAATACGATGCTATGAACCCTTTATTGGGATTCATGGGAACAGTCCCAACATCATCTAAAAGGAGAATTCAATAAGCATTGAACCTCGCCTAAAGACTTTAAGCAAAGCAAAGAGAAAGCGAAGAGCTTTAAAGACAATTGAGAGACAAATCACAAGAATCGAGAGACTTTAA<br>GAGGAAACCAAATTACAAAGAGATGTCTTGGATAAAACCCACTAACCTCACTCCTACCTCGTCGCCCATATTGAGGAAAAGGATAGAGAAATTTTGAAATATAAATCCCTTTTAAAGAAAATGGATTAGGATTATTAGATCCTAAAAGCAGAAAACGATGATGTAGCCAATGTAATTATTAGTGTATAAAT<br>TTAATAGAGATTCATGCGAGCTGAAGAGGATCTATAGAGATTGAGAATTAAAAGACAGAACCTACAAAACATTTAGAACATTTTGATTGGTATTGCCACTGGTAGTTAGAATGATATGAGATAAAATCTGAGAAGTGCCATAAATGAGATGTCGGAAGCCTTAAATACAAAGACATATATTGATCCTCAAA<br>GTGCGATCTAAAAAAAGGGTATGAATTTGCATTAATTGCAAGGTGGAATTCAAAGTAGTTAATAAAAATAGGAAGATGGTCCTTCTTGGTATTCAAAATTGAAACAAAAACAGAATAAAGGAGGAGGACTCGAGGTACGCATGTCTAAAGGAGAAGAATTATTCACTGGTGTTGTTCCAATTCTTGTTGA<br>ACTTGATGGTGATGTTAATGGACATAAATTTTCTGTTTCTGGTGAGGGTGAAGGTGATGCAACATATGGAAAATTAACCTTAAAATTTATTTGCACTACTGGAAAATTACCTGTTCCATGGCCAACACTTGTCACTACTTTAACTTATGGAGTTCAATGTTTTTCAAGATACCCTGATCATATGAAACAACAT<br>GACTTTTTCAAATCTGCCATGCCAGAAGGATATGTCCAAGAAAGAACTATATTCTTCAAAGATGATGGAAACTACAAGACAAGAGCTGAAGTCAAATTTGAAGGAGATACCCTTGTCAATAGAATTGAGTTAAAAGGAATTGATTTTAAAGAAGATGGAAACATTTTAGGACATAAATTGGAATACAACTA<br>TAACTCACATAATGTATACATCATGGCAGACAAACAAAAAAATGGAATCAAAGTCAACTTCAAAATTAGACACAACATTGAAGATGGATCAGTTCAATTAGCAGACCATTATCAACAAAATACTCCAATTGGAGATGGCCCTGTTTTATTACCAGACAACCATTACTTATCAACACAATCTGCCTTATCAAA<br>AGATCCAAATGAAAAGAGAGACCATATGGTTTTATTAGAGTTTGTAACTGCTGCTGGAATTACACATGGCATGGATGA |
| | **siRNA mammalian cells** | |
| **siRNA CEP90 #1** | siGENOME SMARTpool siRNA D-020125-01, PIBF1 Dharmacon | GAGAUUAGACAACCAAUUG |
| | siGENOME SMARTpool siRNA D-020125-02, PIBF1 Dharmacon | ACAGGUAUCUCAUUGAAUC |
| | siGENOME SMARTpool siRNA D-020125-03, PIBF1 Dharmacon | UCGUUAAGAUGCAUAGUAA |
| | siGENOME SMARTpool siRNA D-020125-04, PIBF1 Dharmacon | GAUCAGCUCUUAGACAGGU |
| **siRNA CEP90 #2** | based on siRNA CEP90#1 (Kodani *et al.* 2015) Dharmacon | TCTGCAAGAGGAAACAGCAAGAAAT |
| **siRNA FOPNL #1** | based on siRNA FOR20#2 (Sedjai et al. 2010) Dhamacon | GTATTATAAAGGCCCTTAA |
| **siRNA FOR20 #2** | siGENOME SMARTpool siRNA D-016243-01, FOPNL Dharmacon | GUACUAACAUUGAAGAUCU |
| | siGENOME SMARTpool siRNA D-016243-02, FOPNL Dharmacon | GCCAAUGGAUGACCACCUA |
| | siGENOME SMARTpool siRNA D-016243-03, FOPNL Dharmacon | GAUCCGAGCUGAAGUUUUC |
| | siGENOME SMARTpool siRNA D-016243-04, FOPNL Dhamacon | GCCAUUUCUUGCGUGGAA |
| **scrambled control** | siGENOME Non-Targeting siRNA Pool #2 Dharmacon | UAAGGCUAUGAAGAGAUAC |
| | | AUGUAUUGGCCUGUAUUAG |
| | | AUGAACGUGAAUUGCUCAA |
| | | UGGUUUACAUGUCGACUAA |
